# You can move, but you can’t hide: identification of mobile genetic elements with geNomad

**DOI:** 10.1101/2023.03.05.531206

**Authors:** Antonio Pedro Camargo, Simon Roux, Frederik Schulz, Michal Babinski, Yan Xu, Bin Hu, Patrick S. G. Chain, Stephen Nayfach, Nikos C. Kyrpides

## Abstract

Identifying and characterizing mobile genetic elements (MGEs) in sequencing data is essential for understanding their diversity, ecology, biotechnological applications, and impact on public health. Here, we introduce geNomad, a classification and annotation framework that combines information from gene content and a deep neural network to identify sequences of plasmids and viruses. geNomad uses a large dataset of marker proteins to provide functional gene annotation and taxonomic assignment of viral genomes. Using a conditional random field model, geNomad also detects proviruses integrated into host genomes with high precision. In benchmarks that included diverse MGE and chromosome sequences, geNomad significantly outperformed other tools in all evaluated clades of plasmids and viruses. Leveraging geNomad’s speed and scalability, we were able to process public metagenomes and metatranscriptomes, leading to the discovery of millions of new viruses and plasmids that are available through the IMG/VR and IMG/PR databases. We anticipate that geNomad will enable further advancements in MGE research, and it is available at https://portal.nersc.gov/genomad.

## Main

Mobile genetic elements (MGEs) are selfish genetic entities that, unlike cellular organisms, are unable to self-replicate and instead rely on host cells and cellular machinery to propagate. MGEs are associated with all domains of life and are incredibly diverse, encompassing elements with various replication and mobility strategies, such as plasmids and viruses. Mobile elements are also ubiquitous in nature and found across virtually all of Earth’s ecosystems. Plasmids, for instance, are found in the vast majority of bacterial and archaeal isolates^1^, whereas viruses have been shown to be the most abundant biological entities in oceans and harbor a large reservoir of genetic diversity^2^. Due to their mobility, plasmids and viruses can serve as key drivers of horizontal gene transfer (HGT), a process in which cells acquire genetic information from a mobile gene pool rather than through vertical descent. This process allows distantly related lineages to exchange genetic material, enabling rapid phenotypic shifts that can facilitate adaptation to environmental or biological pressures^3, 4^. As a result, plasmids and viruses play a significant role in driving fast evolutionary and ecological innovation, greatly impacting the dynamics of all biological communities.

With the increased availability of metagenomic sequencing data from diverse ecosystems, it became possible to study the diversity and distribution of MGEs on a global scale. In recent years, numerous studies have harnessed these data to uncover an unprecedented diversity of viral genomes, greatly expanding our understanding of their genetic diversity, distribution, function, and evolution. Plasmids, on the other hand, have been mostly overlooked in metagenomic surveys^5, 6^ and most known sequences are derived from clinical isolates, highlighting the need for further research to understand the factors underlying their spread and evolution in natural environments. Because the HGT promoted by plasmids and viruses can affect the host ecology, metabolism, virulence, and resistance to antibiotics, the identification of these MGEs in sequencing data is also crucial to allow holistic investigation of biological communities. Additionally, detection of MGEs also has important implications for biotechnology and public health, where monitoring virulent strains or sequences carrying antibiotic resistance genes can help prevent the spread of diseases.

Computational identification of plasmids and viruses from sequence data relies on the use of sequence classification models, which can be broadly categorized into two types: alignment-free models and gene-based models. Alignment-free models perform classification directly from nucleotide sequences and employ deep-learning architectures such as recurrent neural networks or convolutional neural networks to learn discriminative sequence motifs that are informative for classification^7^. By dispensing explicit alignments to reference genomes or proteins, these models are theoretically not constrained by the sensitivity of sequence search algorithms or the completeness of reference databases, which often lack close homologues for the fast-evolving genes encoded by MGEs. However, alignment-free models are typically uninterpretable and do not capitalize on prior biological knowledge for classification, hence they are prone to produce overly-confident classification mistakes that can be difficult to diagnose (out-of-distribution generalization problem)^8^. In contrast, gene-based classification methods perform database searches and alignments to identify marker proteins that are indicative of the underlying identity of the sequence^9^. As a consequence, these methods leverage existing biological insights and human-designed features, providing more interpretable outputs. Both alignment-free and gene-based approaches have been used successfully for plasmid and virus identification, but no existing tool effectively combines the strengths of both methods in a single framework.

Here we introduce geNomad, a tool for concurrent identification and annotation of both plasmids and viruses in sequencing data. We demonstrate that geNomad’s classification framework, which utilizes a hybrid approach that combines alignment-free and gene-based models, significantly outperforms other plasmid and virus identification tools across various host and virus taxa. A newly assembled set of 227,897 marker protein profiles that is included in geNomad enables the delimitation of integrated viruses (proviruses), taxonomic assignment of viruses, and rich functional annotation, which includes the identification of antimicrobial resistance genes, conjugation genes, and plasmid and virus hallmark genes.

Applying geNomad to metagenomes and metatranscriptomes revealed numerous RNA and giant virus sequences that were missed by large-scale surveys, significantly expanding phylogenetic diversity of giant viruses. Additionally, we show that geNomad is computationally efficient and scalable, making it suitable for use in large-scale surveys, such as identification of potential virus and plasmids sequences across all public genomes and metagenomes in the IMG database^10^.

## Results and discussion

### The geNomad framework for identifying and annotating virus and plasmid sequences

geNomad employs a hybrid approach to plasmid and virus identification that combines an alignment-free classifier (sequence branch) and a gene-based classifier (marker branch) to improve classification performance by capitalizing on the strengths of each classifier. geNomad’s framework consists of five stages (Figure 1A): (1) alignment-free classification in the sequence branch; (2) sequence annotation and gene-based classification in the marker branch; (3) aggregation of the branch scores; (4) score calibration; and (5) output generation.

**Figure 1.**
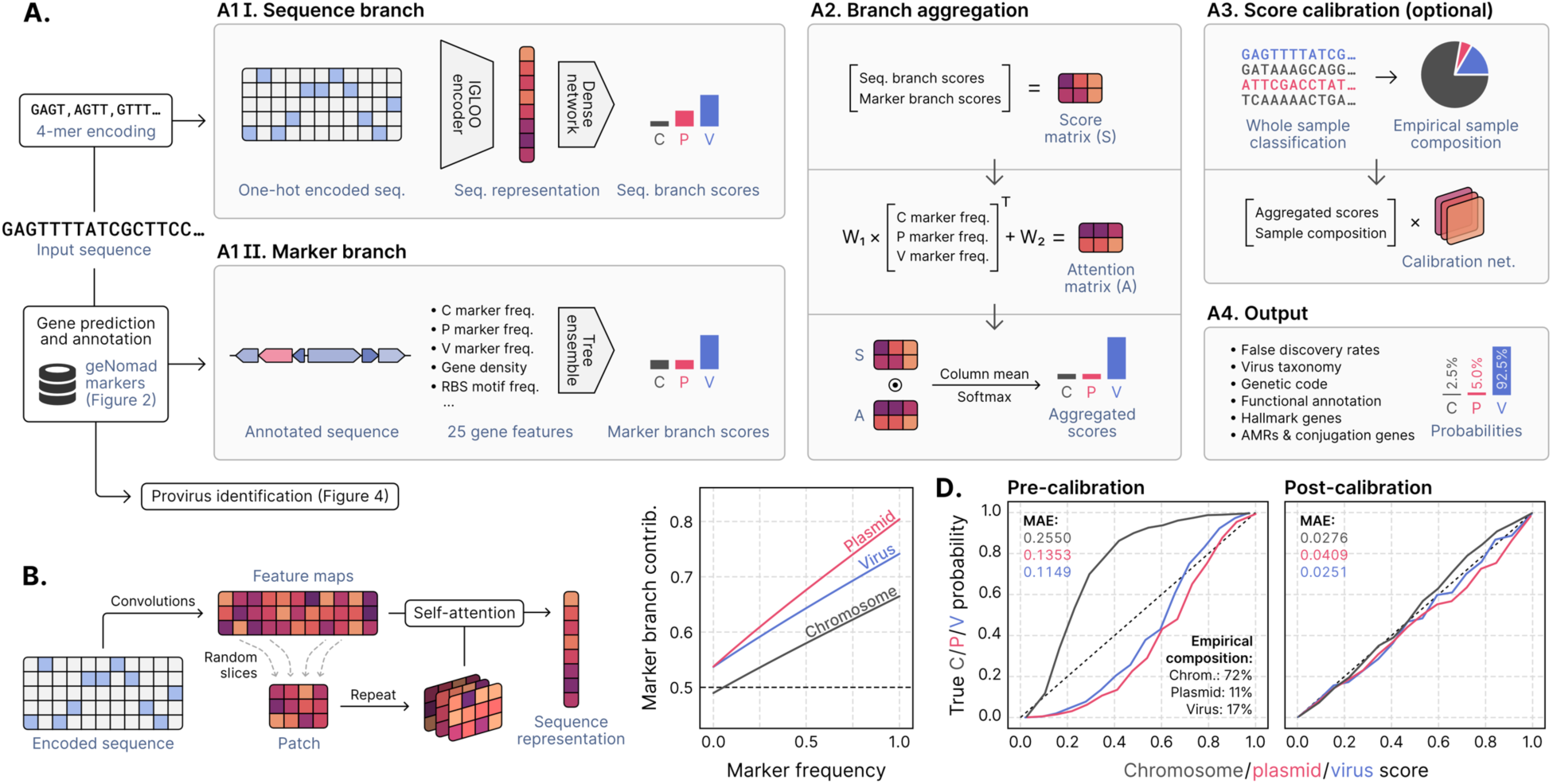
A hybrid framework for identifying and annotating plasmids and viruses. **(A)** geNomad processes user-provided nucleotide sequences through two branches. In the sequence branch, the inputs are one-hot-encoded fed to an IGLOO neural network, which scores inputs based on the detection of non-local sequence motifs (A1 I). In the marker branch, proteins encoded by the input sequences are annotated using markers that are specific to chromosomes, plasmids, or viruses (A1 II). A set of numerical features are then extracted from the annotated proteins and fed to a tree ensemble model, which scores the inputs based on their marker content. Next, the scores provided by both branches are aggregated by weighting the contribution of each branch based on the frequency of markers in the sequence (A2). Aggregated scores can then be calibrated to approximate probabilities in a process that leverages the sample composition inferred from the classification of sequences from the same batch (A3). Lastly, classification results are summarized and presented together with additional data, such as virus taxonomy, gene function, and the inferred genetic code (A4). **(B)** The sequence branch is based on the IGLOO architecture, which uses convolutions to produce a feature map from a one-hot-encoded input. Patches encoding non-local relationships within the sequence are then generated by slicing the feature map. Lastly, these patches are used as an attention matrix to produce a sequence representation from the feature map. **(C)** The relative contribution of the marker branch (*y*-axis, quantified using SHAP) increases as the marker frequency (fraction of genes assigned to a marker) in the sequence increases. **(D)** Calibration curves of pre-calibration (left) and post-calibration (right) scores, showing that sample composition can be used to map classification scores to actual probabilities. The *x*- axis represents scores averaged across multiple bins; the *y*-axis represents the fraction of positives in each bin; the 45° dashed line represents a perfect calibration scenario. MAE: mean absolute error of the scores relative to the true probabilities.

To identify sequences of plasmids and viruses in an alignment-free manner, geNomad’s sequence branch processes inputs using a neural network model that is able to classify the sequences from their nucleotide makeup alone (Figure 1A, box A1 I). To accomplish this, the input sequences are first converted to a numerical format by tokenizing them into arrays of 4-mer words, which are then one-hot-encoded, creating binary 256-dimensional matrices that reflect the presence of specific 4-mers (rows) across different positions within the sequence (columns). These matrices are then passed to an encoder, which generates vector representations of the sequences in a 512-dimensional embedding space. In this space, sequence representations from the same class (chromosome, plasmid, or virus) will be more similar compared to sequence representations from different classes. The resulting representations are subsequently fed to a dense neural network that produces three scores, representing the model’s confidence of the sequence belonging to each of the three classes.

To generate vector representations of the inputs, geNomad employs an encoder based on the IGLOO architecture^11^, which is able to extract patterns that are useful for classification from the sequence data and encode them into an embedding space (Figure 1B, Supplementary Figure 1). The IGLOO encoder begins processing one-hot-encoded matrices by applying 128 convolutional filters to generate sequence feature maps. To gather relationships between non-contiguous parts of the sequence, IGLOO generates 2,100 patches, each containing slices extracted from random positions within the sequence. These patches are subsequently integrated in a self-attention mechanism, where different parts of the feature map are weighted, leveraging the long-range dependencies encoded in the patches, to derive the final sequence representation. The IGLOO architecture has demonstrated superior performance compared to traditional alternatives (such as recurrent neural networks and convolutional neural networks) when applied to sequence data. This is attributed to its capability to gather information from non-local relationships across the entire sequence to create a global representation^11, 12^.

To classify sequences based on their gene content, geNomad’s marker branch predicts and annotates the proteins encoded by input sequences using a set of custom markers that are informative for classification (Figure 1A, box A1 II). To predict proteins, geNomad uses a modified version of the Prodigal^13^ software called prodigal-gv, which we developed to allow automatic detection of TAG-recoded stop codons (common in phages of the *Crassvirales* order^14^) and annotation of TATATA motifs that are frequently found upstream of coding sequences of *Nucleocytoviricota* viruses^15^ and which facilitate their identification. Predicted proteins are then queried against a set of 227,897 protein profiles — specific to chromosomes, plasmids, or viruses (Figure 2) — using MMseqs2^16^ protein-profile search. Next, geNomad computes a total of 25 numeric genomic features that summarize the sequence structure (e.g., gene density, and strand switch rate), RBS motifs (e.g., TATATA motif frequency), and marker content (e.g., frequency of chromosome, plasmid, and virus markers) of the input sequences. These features are then fed to a tree ensemble classification model, which outputs the confidence scores for each class. Detailed explanations of the features used for classification can be found in the Supplementary Note and their importance for the model is shown in Supplementary Table 1.

**Figure 2.**
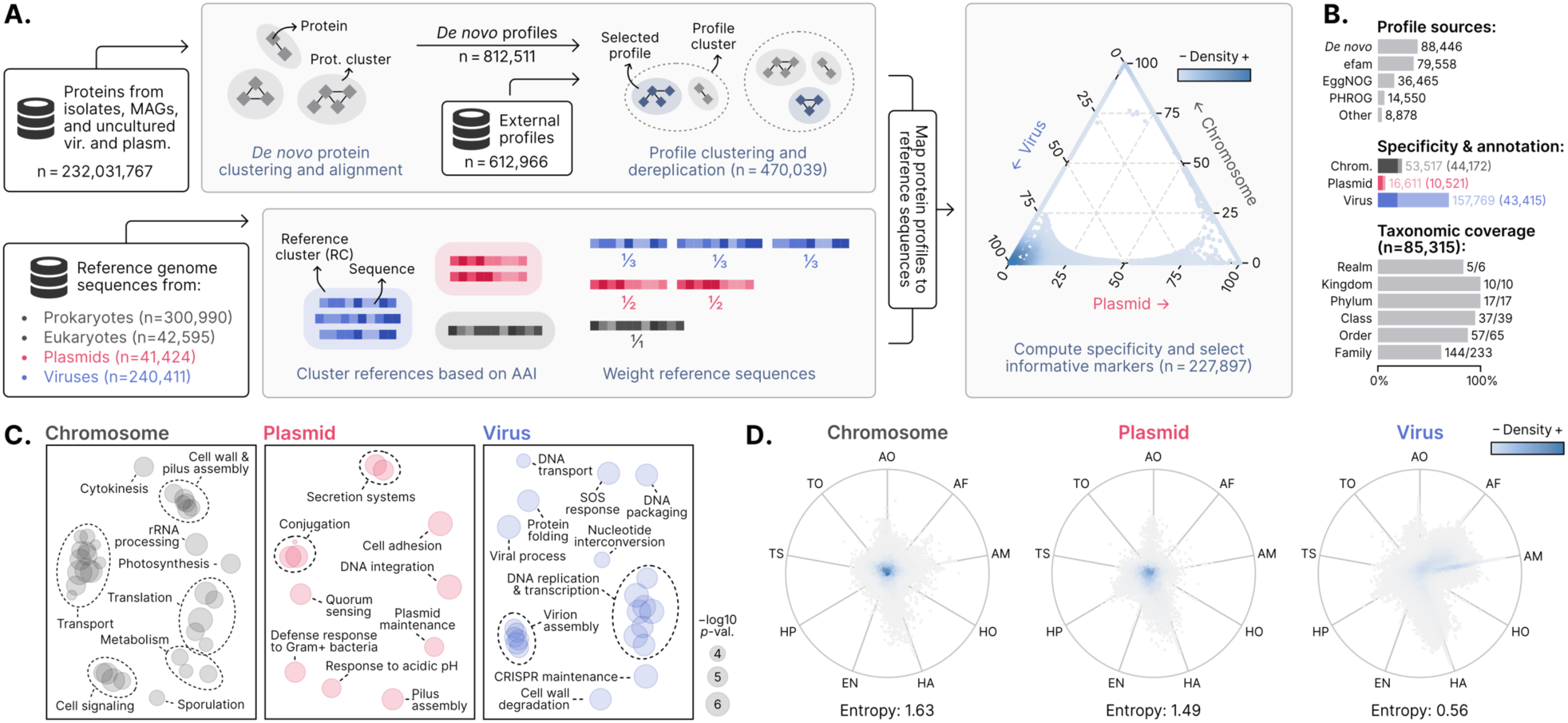
Generating of a dataset of protein profiles with abundant metadata for sequence classification and protein annotation. **(A)** Protein sequences from genomes and metagenomes were clustered and aligned to produce *de novo* protein profiles. *De novo* profiles and profiles obtained from public databases were then clustered and cluster representatives were selected to reduce redundancy. In parallel, reference chromosome, plasmid, and virus sequences were clustered into reference clusters (RCs). Sequences were then weighted in such a way that the sum of the weights within each RC was constant. Representative protein profiles were mapped to reference sequences and chromosome-, plasmid-, and virus-specificity metrics were computed for each profile based on the weighted number of hits to sequences of each class. Markers that were highly specific to one of the three classes were then selected. The position of each selected marker (circles) in the ternary plot is determined by its specificity, and the colors represent the marker density in a region. **(B)** Bar plots showing: the sources of the selected profiles (upper plot); the total number of markers (light shades) and the number of functionally annotated markers (dark shades) for each class (middle plot); the fraction of ICTV taxa covered by the taxonomically-informative markers at each rank. **(C)** Multidimensional scaling of semantic similarities of the GO terms enriched in chromosome (left), plasmid (center), and virus (right) markers. Labels of related terms were aggregated for clarity. Semantic similarities were computed with REVIGO. **(D)** RadViz visualizations of the relative frequencies of geNomad markers across distinct ecosystems. Each marker is represented by a circle and the colors depict the marker density within a region. The position of the markers in the plot is determined by their frequency in each environment. Markers close to the center of the plot were found in similar frequencies across all ecosystems. Median entropies of the ecosystem distributions are shown below the plots. AO: aquatic (other), AF: aquatic (freshwater), AM: aquatic (marine), HO: host-associated (other), HA: host-associated (animals), EN: engineered, HP: host-associated (plants), TS: terrestrial (soil), TO: terrestrial (other).

From the outputs produced by the sequence and marker branches, geNomad generates an aggregated classification that leverages the strengths of each approach, as the two approaches use distinct and often complementary methods to classify input sequences. Aggregated classification is achieved through an attention mechanism that consists of a linear model that weighs the branches based on the frequency of chromosome, plasmid, and virus markers in the input sequence (Figure 1A, box A2). The attention mechanism works in such a way that the contribution of the marker branch goes higher as the fraction of genes that are assigned to markers increases (Figure 1C). Essentially, the branch aggregation gives more weight to the marker branch when it is more informative (i.e., when most of the genes encoded by the input sequence are assigned to markers) and relies more on the sequence branch when marker information is scarce. This allows geNomad to take advantage of both marker-based and alignment-free classification approaches in a principled manner.

During inference, a classification model assigns a score to each prediction, indicating the level of confidence in that prediction, with higher values representing more confident predictions. However, these scores do not reflect the true probabilities of the predictions being correct, as classification models will exhibit varying false discovery rates when classifying samples with distinct underlying composition. For example, if the same classification model is used to identify viruses in a metagenome (where cellular sequences outnumber viral sequences) and in a virome (that is enriched in viral sequences), it is expected that the model will yield a higher proportion of false positive viruses in the metagenome, where more cellular sequences (that are prone to be misclassified as viruses) will be present (Supplementary Figure 2A). The cause of this issue is that models assign the same score to a given sequence regardless of the composition of the rest of the sample. To address this, we devised an optional calibration mechanism in geNomad that leverages sample composition data to approximate the true underlying probabilities. The algorithm consists of a dense neural network that takes raw scores and the empirical sample composition (i.e., the frequency of chromosomes, plasmids, and viruses in the pre-calibration classification) as inputs and outputs calibrated scores (Figure 1A, box A3) that accurately approximate probabilities (mean absolute errors for pre- and post-calibration scores in Figure 1D). Because this process depends on reliable estimates of the underlying compositions, it works best for samples with sufficient size (e.g., ≥ 1,000 sequences), for which the mean absolute error of the calibration is very low (≈ 1%, Supplementary Figure 2B). In essence, the calibration mechanism adjusts raw scores by reducing or increasing the scores of a given class (chromosome, plasmid, or virus) when its frequency within the sample is low or high (Supplementary Figure 2C and D). When the sample composition is very uneven, this tends to result in large changes in raw scores, while very high or low scores are less affected (Supplementary Figure 2C and E). The calibrated scores produced by geNomad offer users two benefits: (1) estimated probabilities can be used to compute false discovery rates, allowing users to make more informed decisions (e.g., setting a threshold to achieve a desired proportion of false positives), and (2) improved classification performance by adjusting the assigned labels of some sequences after calibrating scores (for more details see the “*geNomad outperforms other tools for plasmid and virus classification, and allows accurate taxonomic assignment of viruses*” section).

Sequences classified as viral with geNomad’s markers are then assigned to taxa defined by the International Committee on Taxonomy of Viruses (ICTV)^17^. This process is made possible by the fact that more than 85 thousand of the markers are specific to a virus taxon (for more details see the “*A dataset of marker protein profiles with rich functional and taxonomic metadata*” section). Briefly, geNomad first designates a taxon to each gene annotated with a taxonomically-informed marker. Then, weights are computed for each taxon included among the gene-level assignments, as well as their parent taxa (up to the root of the taxonomy) by summing the bitscores obtained from the alignments with marker profiles. Finally, the taxonomy of the sequence is determined as the most specific taxon that is supported by at least 50% of the total weight (sum of the bitscores of all genes with taxonomy, Supplementary Figure 3).

Upon completion of its execution, geNomad produces a list of sequences that have been classified as either plasmids or viruses. This list can be refined using additional user-adjustable filters, such as a minimum score, maximum false discovery rate (if score calibration was performed), or a minimum number of plasmid or virus hallmark genes (that are involved in key plasmid or virus functions). At this stage, sequences encoding multiple universal single-copy genes, which are rarely found in MGEs, can also be excluded. The generated output includes rich metadata that can be useful for downstream analysis (Figure 1A, box A4), including model scores (uncalibrated or calibrated), predicted genetic code (as inferred by prodigal-gv), structural and functional gene annotation, presence of conjugation and antimicrobial resistance (AMR) genes, number of hallmark genes, and virus taxonomy. The user is also provided with the nucleotide and amino acid sequences of the identified plasmids and viruses.

### A dataset of marker protein profiles with rich functional and taxonomic metadata

geNomad uses a marker set of 227,897 protein profiles specific to chromosomes, plasmids, or viruses to perform classification based on gene content and to provide functional information for processed sequences (Figure 2A). To build this marker dataset, which covers sequences from uncultured microorganisms and viruses from diverse environments, we gathered approximately 232 million protein sequences from diverse isolates (bacteria, archaea, and viruses), metagenome-assembled genomes (MAGs, from bacteria and archaea), and uncultured viruses and plasmids from diverse ecosystems (see complete list of sources in the “*Database of chromosome, plasmid, and viral sequences for training and benchmarking*” section). These sequences were then clustered and the resulting protein clusters were independently aligned to generate 812,511 *de novo* protein profiles, which can be used to perform sensitive searches for homologs. We supplemented these *de novo* profiles with 612,966 additional protein profiles from external sources and performed dereplication by clustering similar profiles and selecting a representative for each cluster, resulting in a non-redundant set of 470,039 profiles (Supplementary Figure 4A). Compared to other sources, *de novo* profiles comprised the majority of the diversity of profile clusters, as 44.0% of these clusters contained at least one *de novo* profile and 29.9% of the clusters contained only *de novo* profiles (Supplementary Figure 4B). This dereplication process was carried out to obtain a reduced set of profiles that cover as much of the protein sequence space as possible while improving geNomad’s computational efficiency by reducing the time required for sequence searches.

To identify profiles that are informative for classification, we computed the specificity of each profile to each one of the targeted classes (chromosomes, plasmids, and viruses) by mapping them to reference genomic sequences (Supplementary Figure 4C). This diverse dataset of references was obtained by retrieving both isolate and uncultivated (bacteria, archaea and eukaryote MAGs, and uncultured viruses and plasmids) nucleotide sequences and curating the data to remove sequences that would lead to misrepresentations of the underlying specificity of each profile (such as phages integrated in host genomes, viruses and chromosomes mislabeled as plasmids, and plasmid and virus scaffolds binned within MAGs). To measure the specificity of the profiles, we matched them to reference protein sequences and counted the number of hits in each targeted class in a weighted manner, taking into account representation bias. Because plasmid and virus sequences in public databases are heavily skewed towards elements that infect a limited range of microbes (such as model species, pathogens, and human-associated microbes), we downweighted sequences belonging to overrepresented taxa by grouping them into reference clusters (RCs) based on high average amino acid identity (AAI). We assigned weights to the references so that the sum of the weights in all RCs was constant, effectively downweighting sequences within large RCs and preventing a few taxa from dominating the marker selection process^6^. After computing specificity, we discarded profiles that were poorly specific or matched few proteins, resulting in a final set of 227,897 profiles to be used for classification and annotation. Most of the markers originated from the *de novo* protein profile dataset (38.8%), the efam^18^ database of protein families from uncultivated viruses (34.9%), and EggNOG^19^ (16.0%) (Figure 2B, top, Supplementary Table 2). Virus-specific markers dominate the dataset (69.2%), followed by chromosome-specific (23.5%), and plasmid-specific markers (7.3%) (Figure 2B, middle, lighter shades).

geNomad also provides users with detailed taxonomic and functional information to facilitate biological interpretation of the results, enabling more thorough analysis of the identified sequences. To allow this, geNomad markers were functionally annotated using sensitive HMM-HMM alignments against databases of protein families with well-annotated functions (Pfam-A^20^, TIGRFAM^21^, KEGG Orthology^22^, and COG^23^). A total of 98,108 (43.0%) markers were annotated, though the proportion of annotated markers varied significantly among the different specificity classes, with chromosome-specific markers having the highest annotation rate (82.5%), followed by plasmid-specific markers (63.3%), and virus-specific markers (27.5%) (Figure 2B, middle, darker shades, Supplementary Table 2), highlighting the limited availability of functional information for viral proteins. Functional enrichment analysis of the annotated markers (Figure 2C) showed that chromosome markers were enriched in functions related to translation, transport, and metabolism; plasmid markers were enriched in quorum sensing and motility functions; and virus markers were enriched in functions related to virus replication and assembly. To identify hallmark markers — which can serve as strong evidence of a given sequence being classified as a plasmid or virus — we manually selected 967 plasmid and 14,626 virus markers annotated with functions directly related to core functions of these MGEs (such as conjugation genes for plasmids and capsid proteins for viruses). To provide additional context for MGE research, markers were also annotated using specialized databases for specific domains of interest (Supplementary Table 2). This resulted in the identification of 481 markers for genes involved in conjugation and 382 markers for antimicrobial resistance, annotated through alignment with the CONJscan^24^ and NCBIfam-AMRFinder^25^ databases, respectively. In addition, 741 markers for universal single-copy genes (USCGs), which are infrequently present in MGEs and can help reduce false positives, were identified through comparison with profiles from the BUSCO dataset^26^.

To allow taxonomic assignment of viruses using geNomad’s markers, virus taxa from the ICTV (Virus Metadata Resource version 19) were assigned to 85,315 markers. This was accomplished by aligning the markers to viral proteins from NCBI’s NR and then obtaining a consensus taxonomy for each one based on the taxonomy of its hits (details in the *Methods* section). The taxonomically-annotated markers can be used to assign virus sequences to a significant fraction of the viral taxa up to the family rank (Figure 2B, bottom), as at least one marker was assigned to 83.3% of the realms (the only realm missing is *Ribozyviria*), 100% of the kingdoms and phyla, 94.9% of the classes, 87.7% of the orders, and 61.8% of the families. Most of these markers were assigned to the *Caudoviricetes* class (93.1%), which dominates metagenomic data^10^, but other major taxa, such as *Riboviria* (2.8%), *Nucleocytoviricota* (2.2%), and *Monodnaviria* (0.7%), are also largely covered (Supplementary Table 2).

Our marker selection process was designed to maximize the range of covered uncultivated genomes found globally. To evaluate the environmental breadth of geNomad’s markers, we used them to scan a total of 2.3 billion proteins from 28,865 metagenomes and 7,258 metatranscriptomes of various ecosystems. For each marker class (chromosome, plasmid, and virus-specific), frequencies in each ecosystem were used to build RadViz visualizations of the environmental distributions of the markers (Figure 2D). This revealed that while chromosome and plasmid-specific markers are generally not specific to any particular environment (high density near the center of the RadViz visualization and high average entropy of frequencies), virus-specific markers tend to be found in restricted ecosystems (low density over the entire RadViz visualization and low average entropy of frequencies). This suggests that the gene repertoire of uncultivated viruses is highly variable and highlights the importance of incorporating environmental data to develop a framework that can cover a large fraction of the virosphere.

### geNomad outperforms other tools for plasmid and virus classification, and allows accurate taxonomic assignment of viruses

To evaluate the classification performance of geNomad and compare it to other virus and plasmid identification tools that use different approaches for sequence classification (Table 1), we used test datasets consisting of diverse sequence fragments with varying lengths (Supplementary Figure 5A). To minimize overestimation of geNomad’s performance due to the presence of similar sequences in the train and test data, we randomly assigned RCs to five different data splits and performed cross-validation using the leave-one-group-out strategy (more details in the *Methods* section), which forced sequences from the same RC to remain together in either the train or test sets. Performance metrics for all tools were measured five times, using each RC as the test set at a time. The following metrics were computed: precision (fraction of true plasmids/viruses among the sequences classified as plasmid/virus); sensitivity (fraction of the true plasmids/viruses that were classified as such); Matthews correlation coefficient (MCC, correlation between the true and predicted labels); and F1-score (harmonic mean of recall and precision).

**Table 1.** Classification methodology and average runtimes of plasmid and virus identification tools. Runtimes were measured across five executions using the hyperfine tool. A random selection of 10,000 metagenomic scaffolds (total of 18.3 Mb, IMG/M Taxon Object ID: 3300038405) was used as input for all tools. All speed measurements were performed in an Amazon EC2 instance (c5.12xlarge, SSD storage). SEM: standard error of the mean; CNN: convolutional neural network; LSTM: long short-term memory neural network.

| Tool | Method | Plasmid | Virus/Provirus | Runtime ± SEM (s) |
| --- | --- | --- | --- | --- |
| geNomad | Hybrid | ✓ | ✓/✓ | 241.73 ± 0.18 |
| geNomad (marker branch) | Marker-based | ✓ | ✓/✓ | 119.20 ± 0.12 |
| geNomad (sequence branch) | Alignment-free (IGLOO) | ✓ | ✓/× | 118.58 ± 0.10 |
| DeepMicrobeFinder | Alignment-free (CNN) | ✓ | ✓/× | 256.03 ± 0.30 |
| DeepVirFinder | Alignment-free (CNN) | × | ✓/× | 710.00 ± 1.75 |
| Phigaro | Marker-based | × | ×/✓ | 1,585.61 ± 3.23 |
| PlasClass | Alignment-free (k-mer freq.) | ✓ | ×/× | 20.50 ± 0.09 |
| PlasX | Marker-based | ✓ | ×/× | 1,965.15 ± 1.02 |
| PPR-Meta | Alignment-free (CNN) | ✓ | ✓/× | 374.41 ± 1.21 |
| Seeker | Alignment-free (LSTM) | × | ✓/× | 758.85 ± 2.93 |
| VIBRANT | Marker-based | × | ✓/✓ | 662.35 ± 9.39 |
| viralVerify | Marker-based | ✓ | ✓/× | 1,641.64 ± 8.12 |
| VirSorter2 | Marker-based | × | ✓/✓ | 6,303.27 ± 4.25 |
| VirSorter2 (all models) | Marker-based | × | ✓/✓ | 6,745.52 ± 10.14 |

By inspecting the classification performance as a function of the similarity to the train data, we found that performance dropped amongst sequences that were more divergent from the train data. However, geNomad still performs rather well on unseen sequences (Supplementary Figure 5B), especially viruses, illustrating its potential for the discovery of new viral taxa. Measurement of geNomad’s performance on sequences with varying marker coverage (i.e., fraction of proteins assigned to markers) revealed that even those that were targeted by no or few markers were still detected due to the sequence branch of the algorithm (Supplementary Figure 5C). When compared to other tools, geNomad presented superior overall classification performance across all sequence length ranges in both plasmid and virus classification tasks (Figure 3A and B, Supplementary Tables 3 and 4). The difference in performance was especially apparent in short sequences (< 6 kb): while the performance of most tools significantly declined due to the limited genetic information in such sequences, geNomad leveraged its extensive marker dataset and alignment-free neural network to extract as much information as possible and maintain high sensitivity and precision. This highlights the usefulness of geNomad in metagenomic and metatranscriptomic assemblies, where most scaffolds are short.

**Figure 3.**
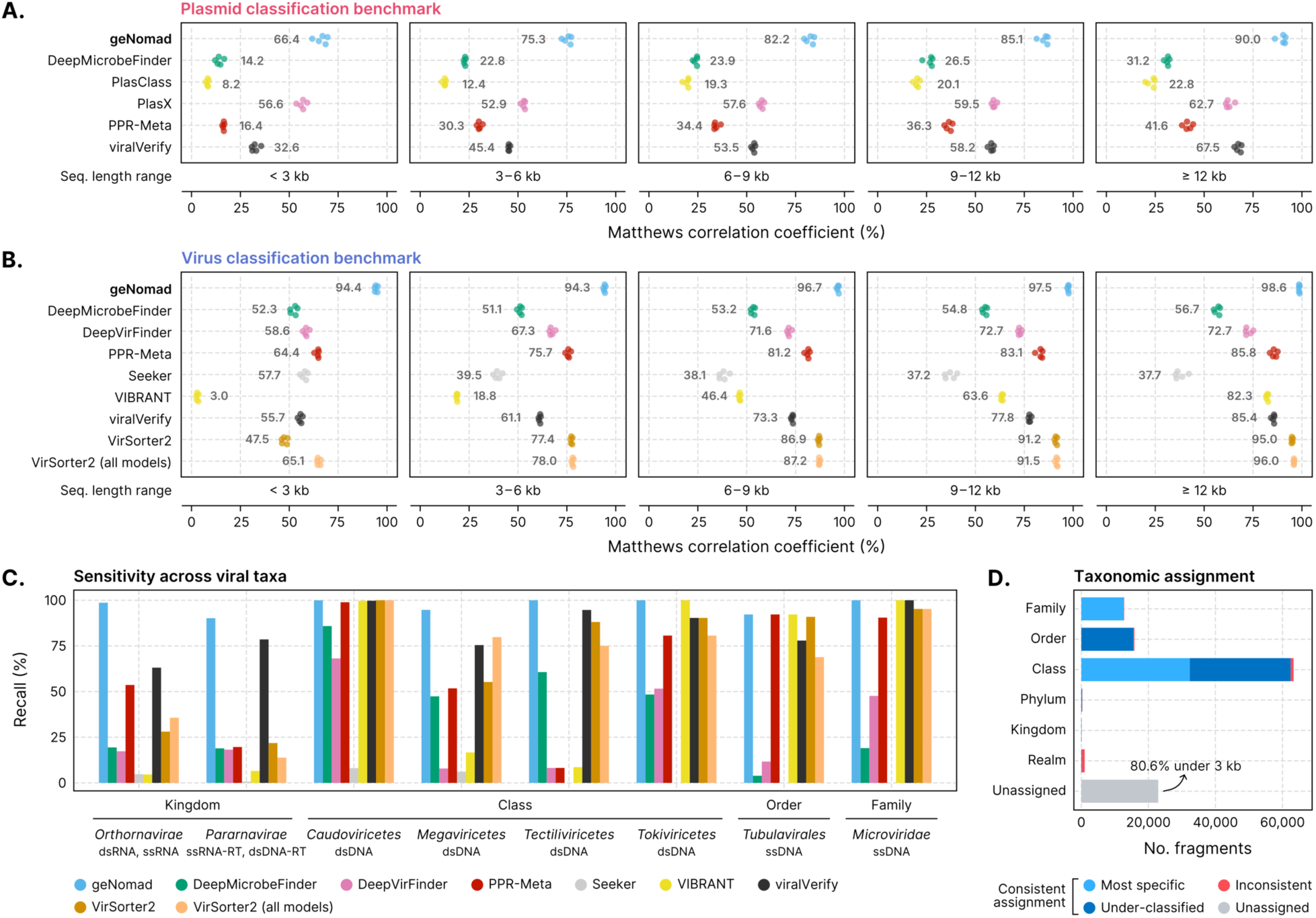
geNomad accurately identifies viruses and plasmids, and allows taxonomic assignment of viral genomes. **(A, B)** Classification performance of multiple plasmid **(A)** and virus **(B)** identification tools across sequence fragments of varying length. Performance was measured using the Matthews correlation coefficient (MCC). For each sequence range interval, tools were evaluated with five different test sets, each containing the sequences of one RC. Coloured circles represent the performances measured in each test set. Mean values are shown next to the circles. **(C)** Sensitivity of virus identification tools across major viral taxa at different ranks. The score cutoff of each tool was determined so that the false discovery rate was approximately 5%. **(D)** Virus taxonomic assignment performance. Bar lengths represent the number of sequence fragments assigned at a given taxonomic rank. Light blue represents sequences that were correctly assigned to their most specific rank (up to the family level); dark blue represents fragments that were assigned to the correct lineage, but to a rank that is above its most specific rank; red represents sequences that were assigned to the wrong lineage; the grey bar represents sequences that were assigned to any taxon.

geNomad’s calibration mechanism enhances the classification process by incorporating sample composition data and assigning estimated probabilities to each sequence, which reflect the likelihood of the sequence belonging to each class. By using calibrated scores instead of raw scores to assign labels, the average classification performance improves because biases introduced during model training are corrected. Indeed, our analysis showed that the plasmid classification performance increased significantly with the use of calibrated scores, particularly for shorter sequences (average ΔMCC: +11.8% for sequences * 3 kb, +5.6% for 3–6 kb, and +3.2% for 6–9 kb) (Supplementary Figure 5D). We also found that short virus sequences benefited from calibration, though the improvement was not as pronounced. These results showcase the effectiveness of the introduced calibration mechanism for improving classification quality.

Plasmid classification is a challenging task due to the variable genetic makeup of these elements, their similarity to other mobile elements that can integrate into host chromosomes, and the lack of a standard for reporting plasmids in sequencing data. As a result, most evaluated tools (DeepMicrobeFinder^27^, PPR-Meta^28^, PlasClass^29^, and viralVerify^30^) had low average classification precision (11.0–40.1%, Supplementary Table 3), even when classifying long sequences (Supplementary Table 4), as they often produced a high number of false positives that can impact downstream analysis. In contrast, PlasX^6^ had high precision (81.6%), but low sensitivity (40.5%), which impairs the detection of plasmids in sequencing data. geNomad had the best overall performance by a significant margin (Figure 3A, MCC and F1-score in Supplementary Tables 3 and 4), with the highest sensitivity (89.8%) and the second highest precision (70.8%), after PlasX. It’s worth noting that geNomad’s marker branch, which can be run independently, achieved a considerably higher precision than PlasX (91.2%).

Most of the plasmid sequences in public databases are limited to a few taxa, such as *Gammaproteobacteria* and *Bacilli*, which can bias the training process if taxonomic imbalance is not taken into account. Because it was designed to reduce the effects of taxonomic representation biases during marker selection and training, geNomad is able to identify plasmids from underrepresented groups more accurately. A similar process was also used in PlasX. When compared to other plasmid identification tools, geNomad had the best performance across all appraised taxa (Supplementary Table 5). Notably, geNomad was the only tool to accurately identify the majority of *Archaea* plasmids (92.54%), which were frequently missed by other tools (3.1–55.3%), and it greatly outperformed other tools for identifying plasmids from major phyla such as *Cyanobacteria* (geNomad: 96.7%, other tools: 6.3–64.3%), *Actinobacteria* (geNomad: 95.5%, other tools: 2.5– 61.9%), and *Bacteroidota* (geNomad: 86.4%, other tools: 2.4–69.2%).

Plasmid identification algorithms can be affected by low quality plasmid annotations in public data. Extrachromosomal viruses and secondary chromosomes are often incorrectly labeled as plasmids in these databases, so it’s important to carefully filter the data to train reliable models and assess classification performance (details in the *Methods* section). To evaluate if existing plasmid identification tools are prone to misclassifying viruses as plasmids — possibly due to contamination in the training data — we measured the fraction of viruses in our test dataset that were labeled as plasmids by the benchmarked tools (Supplementary Table 6). geNomad and PlasX had the best performances in this benchmark (1.7% and 1.5%, respectively), while DeepMicrobeFinder and PlasClass performed the worst (16.0% and 64.4%, respectively). Of note, geNomad’s marker branch classified only 0.2% of the virus sequences as plasmids, which highlights the limitations of current alignment-free tools at this task and the importance of careful dataset curation.

In virus classification, geNomad attained the best overall performance when considering all length strata (MCC: 95.3%, F1-score: 97.3%), followed by VirSorter2^31^ executed with all models (MCC: 81.3%, F1-score: 88.9%), VirSorter2 executed with default parameters (MCC: 79.7%, F1-score: 87.1%), and PPR-Meta (MCC: 77.4%, F1-score: 86.6%) (Figure 3B, Supplementary Table 3). VIBRANT^32^, geNomad, VirSorter2 (default parameters), and viralVerify achieved the highest classification precision (97.5%, 97.3%, 94.7%, and 91.3%, respectively), while Seeker^33^, DeepVirFinder^34^, DeepMicrobeFinder, and PPR-Meta obtained the lowest scores (61.8%, 80.5%, 84.5%, and 88.5%, respectively). VIBRANT’s overall classification performance metrics appeared low (MCC: 36.0%, F1-score: 35.2%) due to its very low sensitivity when classifying short sequences (Figure 3B, Supplementary Table 4), a consequence of it not classifying sequences that encode less than four genes and not being designed to identify eukaryotic viruses (see paragraph below).

The development of tools that can accurately identify diverse viral taxa is challenging, as no genes are universally shared across the virosphere. Additionally, unequal representation of viral groups — illustrated by the dominance of tailed phages from the *Caudiviricetes* class — in sequencing data can bias classification models and prevent the discovery of underrepresented taxa. In a benchmark study using representative genomes from the ICTV, we found that geNomad outperformed other tools in all major taxa we evaluated (Figure 3C, Supplementary Table 7). Notably, geNomad was the only tool that achieved high sensitivity for viruses that encode an RNA-dependent RNA polymerase (*Orthornavirae*, 98.64%), retroviruses (*Pararnavirae*, 90.18%), and giant viruses (*Megaviricetes*, 94.74%) at a fixed false discovery rate of 5%. The only other tool to display sensitivity over 50% for all taxa was viralVerify, while the remaining tools failed to achieve this for at least two of the groups. When evaluating sensitivity across different host clades, we found that geNomad was the only tool that identified more than 90% of the viruses infecting bacteria, archaea, and multiple eukaryotic groups, while other tools struggled to identify viruses that infect at least two of the eukaryotic groups that were evaluated (Supplementary Table 8). In an additional benchmark where we measured classification sensitivity on a catalog of metagenomic *Inovirus*^35^, which are known to be challenging to detect automatically, geNomad (recall: 84.8%) also outperformed other evaluated tools (average recall: 29.9%, Supplementary Table 9). These results show that geNomad can be employed to identify a wide range of virus taxa infecting a variety of hosts, enabling the discovery of viruses that are often missed in metagenomic analyses, such as non-tailed phages and viruses that infect eukaryotes. It is worth noting that several of the tested tools (DeepVirFinder, PPR-Meta, Seeker, and VIBRANT) were trained only on phage data and are therefore not designed to identify viruses that don’t infect prokaryotes. In fact, VIBRANT was a top performer for *Caudoviricetes*, *Tokiviricetes*, *Tubulavirales*, and *Microviridae*.

We assessed the performance of geNomad’s taxonomic assignment (Figure 3D, Supplementary Table 10) by assigning 116,250 artificially fragmented genomes of ICTV exemplar species to viral lineages using a modified marker dataset with modified taxonomic metadata to simulate novelty (see *Methods* for details). Of the processed fragments, the majority (80.3%) was successfully assigned to a viral lineage, with most being classified at the class (54.4%), order (13.6%), or family (10.1%) levels. Among those, 48.2% were correctly assigned to the most specific rank (up to the family level), 49.5% were under-classified (assigned to the correct lineage, but to a rank that is above its most specific rank), and only 2.3% were assigned to the wrong lineage. These results indicate that while geNomad may sometimes under-classify, it is reliable at assigning sequences to higher taxa. The unassigned fragments, which lacked hits to taxonomically-annotated markers, were mostly shorter than 3 kb (80.6%), indicating that geNomad’s marker-based approach will have limited taxonomic assignment sensitivity for short sequences.

### Leveraging geNomad’s markers for sensitive and precise identification of proviruses

Temperate phages can integrate into host genomes and form proviruses, which can greatly affect host metabolism and ecology^36–38^. Several approaches have been developed to demarcate integrated viruses within host sequences, including: identification of virus marker genes^32^, sequence alignment to databases of reference phage genomes^39^, comparative genomics^40^, and mapping of paired-end reads^41^. The sensitivity of methods that rely on comparing sequences to references is limited by the fact that such genomes represent a small fraction of the virosphere. While comparative genomics and read mapping can provide accurate coordinates of the provirus integration sites, they rely on the availability of adequate data, heavy computation, and are unable to detect proviruses that are not active. To achieve high sensitivity and speed, geNomad leverages its extensive marker dataset to identify proviruses based on the annotation of viral hallmarks (Figure 4A).

**Figure 4.**
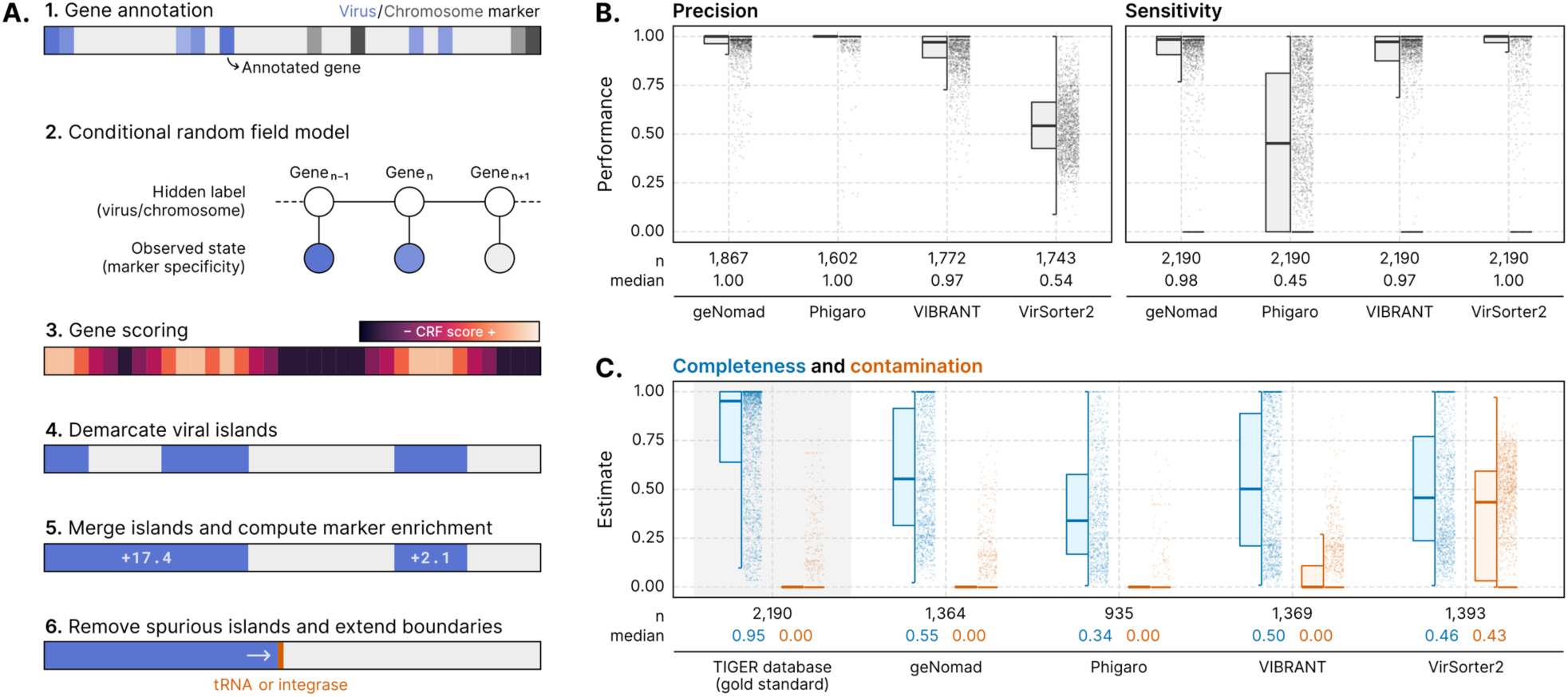
geNomad uses marker information to demarcate provirus boundaries. **(A)** Provirus identification starts by annotating the genes within a sequence with geNomad markers, which store information of how specific they are to hosts or viruses. These specificity values are then fed to a conditional random field model, which will score each gene using information from the markers in its surroundings. A score cutoff is used to demarcate viral islands, and islands that are close together are merged. Islands with few viral markers are discarded and the boundaries of the remaining islands are extended up until nearby tRNAs or integrases. **(B)** Distributions of the precision and sensitivity of multiple provirus-identification tools, measured at the gene-level for each provirus. Proviruses from the TIGER database were used as the ground truth for this benchmark. **(C)** Completeness and contamination estimates of demarcated proviral regions that did not overlap with proviruses in the TIGER database. Estimates for TIGER proviruses are shown with a grey background as a reference.

To demarcate the coordinates of viruses integrated within host genomes, geNomad employs a conditional random field (CRF) model that uses contextual information to search for regions enriched in viral markers that are flanked by chromosome markers. The CRF model used in geNomad takes advantage of the high gene coverage provided by the marker database to score a given gene, taking into account the level of specificity of the markers that were assigned to that gene and its neighbors. To filter out spurious viral islands (regions of consecutive genes that were labeled as viral), geNomad merges islands that are close together and then removes the ones with a low marker enrichment (that is, with few virus markers). Finally, because tRNAs and integrases are commonly found next to the edges of integrated elements due to the dynamics of site-specific recombination^40^, geNomad extends provirus boundaries up until neighboring tRNAs and/or integrases (identified using a supplementary set of 16 site-specific tyrosine integrases), improving the detection sensitivity of genes close to provirus edges, which sometimes include accessory genes, such as those encoding defense systems, that influence the phage-host dynamics^42^.

We assessed geNomad’s provirus demarcation performance and compared it to that of other popular tools (Phigaro^43^, VIBRANT, and VirSorter2) using the TIGER dataset^40^, which contains precisely mapped integration sites across 2,168 prokaryotic genomes, as the ground truth (Figure 4B, Supplementary Table 11). For each proviral region that was predicted by the tools being benchmarked, precision was measured as the fraction of genes that were located within TIGER proviruses. Sensitivity was measured for each TIGER provirus as the proportion of genes that were contained within the regions predicted by each tool. This benchmark revealed that geNomad was able to find more proviruses than the other tools and displayed high precision and sensitivity values. While Phigaro also showed high precision, it failed to detect a large number of proviruses and exhibited very low sensitivity. Conversely, VirSorter2 displayed very low precision — meaning that it left host genes within provirus boundaries — and high sensitivity. Not all of the predicted proviral regions overlapped with TIGER coordinates, since this dataset doesn’t include inactive phages nor proviruses that don’t integrate at tRNAs. To measure the quality of such predictions, we used CheckV^44^ (version 1.0.1) to estimate the quality of each proviral region and found that geNomad outperformed other tools as the proviruses it demarcated tended to be more complete and have low contamination levels (i.e., few host genes) (Figure 4C, Supplementary Table 11). The completeness of most of these proviral regions was, however, notably lower than the proviruses contained in TIGER, indicating that they likely represent inactive proviruses that underwent gene loss.

In an additional benchmark, we compared provirus prediction tools using comparative genomics. We employed PPanGGOLiN^45^ (version 1.2.74) to create a *Pseudomonas aeruginosa* pangenome from 442 genomes and to identify its core genes, which are persistent across genomes and are not expected to be found within proviruses. Next, we measured the fraction of core genes in each predicted provirus region as a proxy for contamination and found that, compared to the other evaluated tools, geNomad retrieved significantly more proviruses that tended to have low contamination levels (Supplementary Figure 6A, Supplementary Table 11). To illustrate the importance of precise boundary demarcation for downstream biological interpretation, we show that geNomad was able to find provirus-encoded defense systems — such as DarTG^46^ and Hachiman^47^, detected with DefenseFinder^48^ (version 1.0.9) — that were missed by overly conservative tools (Phigaro and VIBRANT) while excluding core host genes that were left within prophages by VirSorter2. DarTG was found right next to an integrase, illustrating how leveraging tRNAs and integrases for boundary prediction can improve interpretation of the phage-host interactions (Supplementary Figure 6B).

### geNomad is fast and user-friendly, allowing analysis of large datasets

To enhance user experience and make geNomad accessible to a wide audience, we designed it to be user-friendly and efficient, allowing it to run quickly on a broad range of hardware. geNomad can be installed locally though diverse methods (pip, Conda, and Docker), facilitating its installation in a variety of scenarios. In regards to user experience, we designed an informative command-line interface that contains detailed explanations of the commands and a detailed logging of the executions. To make geNomad more accessible to those who may not be familiar with command-line interfaces, we have also made it available as a web application through the NMDC EDGE platform (available at https://nmdc-edge.org/virus_plasmid/workflow). This allows users to upload their data and visualize the results directly in their web browser. Additionally, the integration with NMDC EDGE enables geNomad to be easily incorporated into larger workflows that include other tasks, such as assembly and binning.

In bioinformatics analysis, execution speed can often be a bottleneck due to limited access to powerful hardware and time constraints faced by researchers. To address this, we aimed to make geNomad as fast as possible through several approaches: (1) employing a dereplication approach to produce a marker dataset that covers as much as possible of the protein space with a reduced number of profiles; (2) utilizing MMseqs2 profile search, which is more sensitive than standard protein searches and is much faster than the commonly used hmmsearch command; and (3) implementing the IGLOO architecture for the sequence branch classifier, rather than traditional architectures like CNNs and LSTMs that are more time-consuming. In a benchmark measuring the time it took to classify 10,000 metagenomic scaffolds (Table 1), geNomad outperformed all but one of the evaluated tools, taking significantly less time than VirSorter2 (26.1× improvement), PlasX (8.1×), viralVerify (6.8×), and VIBRANT (2.7×). The only tool that was faster was PlasClass, which only uses k-mer frequencies for classification and exhibited low classification performance in our benchmarks (Figure 3A). It’s worth noting that geNomad’s marker and sequence branches can be run independently, reducing runtime by half while still maintaining good classification performance (Supplementary Table 3), in cases where time is a concern. These results demonstrate that, due to its speed, geNomad can be used in varied hardware and can be scaled to process large datasets. In fact, geNomad was recently used to process approximately 260 million scaffolds from IMG/M to gather the data used to build the IMG/VR v4^10^ and IMG/PR databases, which represent the largest available databases of virus and plasmid sequences, respectively. In the following sections, we highlight the identification of previously undetected genomes of RNA viruses and giant viruses, two groups that especially benefit from geNomad’s classification framework.

### geNomad allows sensitive detection of RNA viruses in natural environments

Recent studies have revealed a previously undetected diversity of RNA viruses from the *Orthornavirae* kingdom by performing large-scale metatranscriptome surveys^49–51^. However, these surveys are limited by their reliance on detecting the RNA-dependent RNA polymerase (RdRP) hallmark protein, thus systematically overlooking genome segments that do not encode RdRP and fragmented scaffolds missing this gene. As geNomad leverages an extensive set of markers covering diverse functions (1,293 out of the 1,906 markers assigned to *Orthornavirae* are not functionally annotated as RdRP) and an alignment-free classification model that doesn’t rely on gene families, we tested whether it could reliably detect segments or fragmented sequences of RNA viruses that are missing the RdRP protein. To evaluate this, we gathered likely RNA virus sequences that do not encode RdRP by binning metatranscriptomes from microbial communities in the Sand Creek Marshes^52^ based on high read coverage correlation with RdRP-encoding scaffolds. The co-occurrence of a given sequence with another encoding the hallmark protein across multiple samples suggests that they came from the same *Orthornavirae* genome. Importantly, this binning-based approach does not rely on features used by geNomad for classification to identify those scaffolds, allowing us to avoid potential biases in our analysis.

In total, we identified 623 scaffolds that co-occurred with RdRP-encoding sequences across 34 metatranscriptome assemblies. The majority of these scaffolds (98.1%) were classified as viruses, indicating that geNomad is capable of identifying sequences of RNA virus genomes even when they lack the RdRP hallmark protein (Figure 5A). When evaluating how other tools classify these sequences we found that, on average, only 43.3% of the scaffolds were classified as viral and that alignment-free models presented a higher sensitivity (Supplementary Table 12), highlighting that such scaffolds are often not targeted by markers. As expected, sequences containing RdRP genes were almost always classified as viral (99.9%, Figure 5A). Inspection of pairs of co-occurring scaffolds revealed that they fell into two categories: (1) linear genomes that were assembled into two sequence fragments, one of which lacked the RdRP gene (*Marnaviridae* bin in Figure 5B); and (2) segmented genomes, where the genome is encoded across multiple DNA molecules, only one of which encodes the RdRP (*Cystoviridae* bin in Figure 5B). Closer examination of these sequences revealed that they encoded domains associated with viral function, such as helicases, proteases, and structural proteins. Many of these domains were covered by geNomad’s markers (coloured genes in Figure 5B), demonstrating that the use of an extensive set of protein profiles enabled geNomad to sensitively identify fragments of RNA virus genomes. Among sequences not encoding RdRP and not binned with RdRP-encoding scaffolds, yet classified as viruses by geNomad, we found fragments of RNA virus genomes missing the RdRP gene (*Leviviridae* scaffold in Figure 5B) and transcripts of DNA viruses (*Caudoviricetes* scaffold in Figure 5B).

**Figure 5.**
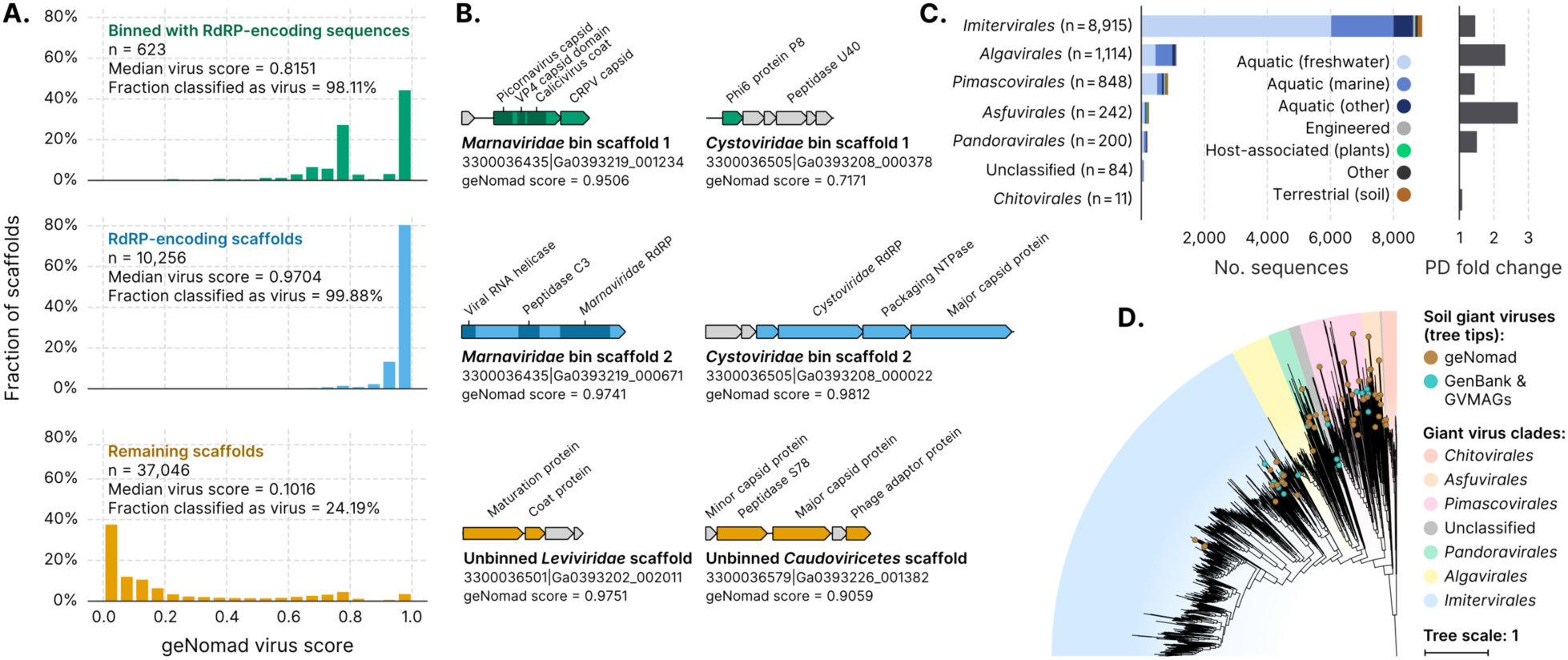
geNomad allows the discovery of RNA viruses and giant viruses in environmental sequencing data. **(A)** Histograms showing the geNomad score distribution of three groups of scaffolds of the Sand Creek Marshes metatranscriptomes: scaffolds that binned with RdRP-encoding sequences (top row, in green); scaffolds that contain the RdRP gene (middle row, in blue); and the remaining scaffolds (bottom row, in orange). The median geNomad score and the fraction of scaffolds classified as viral is indicated for each group. **(B)** Genome maps of selected sequences that were classified as viral by geNomad. Two pairs of co-occurring *Orthornavirae* scaffolds are represented (*Marnaviridae* and *Cystoviridae* bins). Genes targeted by geNomad markers are coloured, while genes that do not match any marker are shown in grey. Rows and colors match those of panel A. **(C)** Number of scaffolds assigned to *Nucleocytoviricota* orders across multiple ecosystems (left bar plot). Sequences were identified by geNomad in a large-scale survey of metagenomes of diverse ecosystems. Only scaffolds that are at least 50 kb long or more were evaluated. Bar colors represent the ecosystem types where the sequences were identified. The phylogenetic diversity (PD) fold change is shown on the right bar plot. PD fold change values correspond to the ratio between the total PD of trees reconstructed with and without geNomad-identified giant viruses. **(D)** Maximum-likelihood phylogenetic tree of soil giant viruses identified with geNomad (brown tree tips). Reference sequences from GenBank and from a previous metagenomic survey (GVMAGs) were included and the ones that were sequenced from soil samples are indicated with turquoise tree tips. Tree tips that are not colored represent representative genomes sequenced from samples obtained from other ecosystems. The ranges corresponding to different *Nucleocytoviricota* orders are represented using distinct colors.

### Application of geNomad in metagenomic data significantly expands the diversity of giant viruses

Giant viruses of the *Nucleocytoviricota* phylum possess large and complex genomes, and their virions can be as large as the cells of many bacteria and archaea^53^. Due to their expansive genomes and diverse genetic repertoires, the identification of these viruses through high-throughput methods is challenging and often relies on computationally expensive phylogenetic analyses and metagenomic binning, which limits the search space^15, 54, 55^. To make geNomad capable of sensitive detection of giant viruses in sequencing data, we expanded the diversity of *Nucleocytoviricota* in the training data by including genomes identified in a previous metagenomic survey (Schulz *et al*. 2020)^15^. Additionally, we included classification features specifically designed to enhance their detection, such as frequency of giant virus-specific markers and the TATATA motifs (Supplementary Table 1, Supplementary Note). As a result, we found that geNomad significantly outperformed other tools in the classification of *Megaviricetes* giant viruses (Figure 3C).

To assess geNomad’s capability to uncover new clades of giant viruses in sequencing data, we applied it to 28,865 metagenome assemblies from the IMG/M^56^ database. Scaffolds classified as virus by geNomad that were at least 50 kb in length were further analyzed using the GVClass pipeline, which placed *Nucleocytoviricota* scaffolds in a phylogenetic context by identifying a set of conserved protein families and reconstructing gene trees together with reference genomes. A total of 11,414 scaffolds identified by geNomad were phylogenetically placed in the *Nucleocytoviricota* tree (median length: 73.3 kb, interquartile range: 58.6–102.7 kb, Figure 5C, Supplementary Table 13). Other tools classified, on average, 71.2% of these scaffolds as viral (Supplementary Table 14). To compare the results with those obtained using the pipeline described in Schulz *et al*. (2020), we examined metagenomes that were processed using both methodologies and found that 1,562 sequences (43% of total) were only detected by geNomad, 1,976 scaffolds (55%) were identified by both methodologies, and only 74 (2%) were found exclusively in the previous survey, demonstrating that geNomad allowed increased recovery of *Nucleocytoviricota* sequences.

The majority of the giant virus sequences identified by geNomad were found to belong to the *Mesomimiviridae* family (n = 6,372) of the *Imitervirales* order (n = 8,915), which includes viruses of haptophytes and ochrophytes^57^ (Figure 5C, Supplementary Table 13). By measuring the increase in phylogenetic diversity brought by scaffolds from this survey, we found that the diversity of multiple orders was significantly expanded (Figure 5C, Supplementary Table 13), particularly that of *Asfuvirales* (2.7× increase) and *Algavirales* (2.3× increase). Within metagenomes from soils, an understudied niche for giant viruses^58^, we identified 235 additional *Nucleocytoviricota* scaffolds, up from 16 metagenomic bins reported in the previous survey. Phylogenetic reconstruction of these soil giant viruses revealed that they include several novel clades of *Imitervirales*, *Pimascovirales*, and *Asfuvirales* that do not have representatives in GenBank or Schulz *et al*. (2020) (Figure 5D), suggesting that the underlying diversity of *Nucleocytoviricota* in soil is greatly underestimated.

## Conclusion

Identifying plasmids and viruses in sequencing data is a crucial process, as it sheds light on the diversity of these mobile elements, their impact on the evolution and ecological interactions of cellular organisms, and facilitates high-throughput monitoring of clinically-relevant strains. Here, we present geNomad, a novel computational framework that enables the identification, annotation, and classification of plasmids and viruses in sequencing data. This is supported by a database of marker protein profiles that are richly annotated in terms of functional and taxonomic information. As a result, this framework has broad application for sequence classification and annotation, allowing, for example, end-to-end identification of conjugative plasmids that carry AMR genes. geNomad incorporates innovative concepts such as a hybrid classification process that combines alignment-free and gene-based approaches in a principled manner, and a score calibration algorithm that enhances the quality and interpretability of results and can be applied in various computational biology applications relying on machine learning. Given its improved classification performance and computational efficiency compared to other tools, as well as its ability to taxonomically classify viruses and functionally annotate genes, we anticipate geNomad will be a valuable resource for the plasmid and virus research communities. We also foresee that it will drive further exploration of the virosphere and foster new initiatives to uncover the diversity and ecology of plasmids in natural environments, an area that has often been overlooked.

While geNomad provides significant performance improvements compared to other tools, downstream quality assessment of the predictions is highly recommended as automated sequence classification algorithms are inherently susceptible to false positives and benchmarks cannot account for all the diversity present in real metagenomics samples. For instance, fragments of genomic islands, transposons, and other mobile genetic elements can have gene content similar to some plasmids, leading to incorrect classification. Manual or automated^44^ quality assessment of virus predictions are also important to identify virus-derived elements, such as gene transfer agents^59^ and tailocins^60^, that are susceptible to be flagged as proviral regions by geNomad since the algorithm does not make a distinction between active and domesticated phages. Therefore, acknowledging such limitations, geNomad has been designed to provide users with abundant information on top of classification results, allowing them to analyze data based on their own criteria.

In the future, geNomad can be improved by incorporating new knowledge on plasmid and virus diversity and biology. As more sequence data become available, new classification models can be trained, and markers can be added or removed to improve the classification performance of new and existing groups of plasmids and viruses. Additionally, new features can be incorporated to provide a more comprehensive toolkit for mobile genetic element (MGE) discovery and annotation. First, binning data could be leveraged to increase the amount of information available for a given genome during classification, an approach that has been shown to improve the classification of phages^61^. This would be particularly useful for giant viruses, which commonly require metagenomic binning as their genomes are rarely recovered on a single scaffold due to their relatively large genome sizes^62^. Second, geNomad could benefit from read mapping to improve the precision of provirus delimitation^41^ and to allow detection of integrative and conjugative elements, which, despite their similarities with plasmids, are frequently overlooked in metagenomic studies as current identification methods rely on comparative genomics^40, 63^. Finally, additional models could build upon geNomad’s markers to further annotate viruses and plasmids, for example, by classifying phages based on their lifestyle (temperate or virulent) and plasmids based on their motility (conjugative, mobilizable, or non-mobilizable), providing users with additional biological information about the identified MGEs.

## Methods

### Database of chromosome, plasmid, and viral sequences for training and benchmarking

Prokaryotic genomes (2,886 bacterial and 336 archaeal) were retrieved from GTDB^64^ (release 202). To reduce taxonomic bias and to limit computational overhead, we only used the genome with the highest quality score (completeness − 5 × contamination − 0.05 × no. scaffolds) per GTDB family. Provirus and provirus-like regions were identified using VirSorter2 (version 2.2.2), Phigaro (version 2.2.6), and VIBRANT (version 1.2.1) and removed from the scaffolds. Plasmids were removed by identifying sequences that either had the word “plasmid” in their header or that shared at least half of their genes with any plasmid in the PLSDB database^65^ (release 2020_11_19). Eukaryotic genomes were obtained from the TOPAZ dataset^66^, which is composed of 988 metagenome-assembled genomes of small eukaryotes that are more likely to be found in metagenomic assemblies. To reduce taxonomic imbalance, we clustered these genomes based on their AAI into 385 clusters using the Leiden algorithm^67^ (as implemented in the igraph Python package, resolution parameter = 0.5) and picked the genome with the least contamination (as estimated by the study’s authors) as the representative.

Plasmid sequences were obtained from the PLSDB database (release 2020_11_19), RefSeq (archaeal plasmids, retrieved in 2021/07/23), and a dataset of complete plasmids identified in metagenomic data (IMG/M Taxon Object ID: 3300053491). To identify chromosome sequences that were mislabeled as plasmids, we performed gene prediction with Prodigal (version 2.6.3, parameters: “-m -p meta”) and used hmmsearch^68^ (HMMER version 3.3.2, parameter: “--cut_ga”) to match the proteins to sets of essential single-copy genes (ar122 and bac120, used for phylogenetic reconstruction in GTDB). Scaffolds encoding two or more essential genes were discarded.

To further remove viral scaffolds from the prokaryotic and eukaryotic chromosome datasets, as well as phage-plasmids from the plasmid data, we performed an additional filter using HMMs of viral hallmarks from VirSorter2 and viral and host markers from CheckV (database version 1.0). Briefly, we used hmmsearch (parameter: “-E 1e-5”) to match Prodigal-predicted proteins from all chromosome and plasmid scaffolds to these HMMs and discarded the sequences that encoded any viral hallmark or that had no. viral markers ≥ 0.5 × no. host markers.

The virus sequence dataset was assembled using data from GenBank (retrieved on 2021/07/06), IMG/VR v3^69^, *Nucleocytoviricota* from Schulz, F. *et al.* (2020)^15^, *Leviviridae* from Callanan, J., *et al*. (2020)^70^, Asgard archaea viruses from Medvedeva, S. *et al.* (2022)^71^, archaeal tailed viruses from Liu, Y. *et al.* (2021)^72^, and *Orthornavirae* from Neri, U. *et al.* (2022)^51^. To remove short genome fragments and contaminants from the IMG/VR sequences, we only retained sequences that contained direct terminal repeats or that fulfilled the requirements to be considered high-quality according to the MIUViG standard^73^. Because the *Nucleocytoviricota* genomes from Schulz, F. *et al.* consist of metagenomic bins that might contain contamination, we opted to keep only the contigs that encode the major capsid protein (MCP), identified using hmmsearch (parameter: “-E 1e-5”) to match their proteins to the set of MCP HMMs provided in the original study.

To reduce sequence redundancy, plasmid and virus scaffolds were de-replicated using pairwise average nucleotide identities (ANI), computed as described in Nayfach *et al.* (2021)^44^ (code available at https://bitbucket.org/berkeleylab/checkv/src/master/scripts/anicalc.py). Specifically, we used Megablast^74^ (version 2.11.0+) to perform all-versus-all nucleotide alignments and computed the pairwise ANI as the length-weighted average identity of all the matches between a pair of sequences. Next, scaffolds with ANI ≥ 97% over at least 95% of the length of the shorter sequence were clustered using a greedy algorithm, as previously described^75^, and the longest sequence within each cluster was selected as the representative. Scaffolds shorter than 2,000 bp were discarded. In total, the final selection contained 300,990 sequences from prokaryotic chromosomes, 42,595 sequences from eukaryotic chromosomes, 41,424 plasmid sequences, and 240,411 virus sequences.

To account for the taxonomic representation imbalance of public databases, plasmid and virus sequences were structured into reference clusters (RCs) containing related sequences. The RCs would serve two main purposes: (1) to minimize representation bias in model training, by downweighting the sequences within large RCs to that the total weight of each RC was the same; (2) to allow us to perform an informed cross-validation splits^6^, where the sequences of a given RC will remain together in either the train or test sets, allowing us to measure geNomad’s performance on novel sequences. To obtain the RCs, we computed the AAI between all pairs of plasmids and viruses and built a network using these values as edge weights (code available at https://github.com/apcamargo/bioinformatics-snakemake-pipelines/tree/main/contig-aai-pipeline). Next, we employed the Leiden algorithm to cluster the scaffolds, tuning the resolution parameter to make the average within-cluster AAI as close to 95% as possible. In total, we obtained 32,134 plasmid RCs and 215,618 virus RCs. Because prokaryotic and eukaryotic scaffolds are organized in genomes, we opted to treat all the sequences within a given genome as members of an effective RC. The RCs were randomly assigned to five distinct data splits that would be used for benchmarking.

Given that metagenomic assemblies are mostly comprised of short sequence segments, we created a dataset of artificially fragmented sequences that would be used for model training and evaluation. We first built an empirical length distribution from all public IMG/M metagenomes (as of 2021/09/11) and truncated the distribution to a minimum of 3,000 bp. Next, we split the sequences of our final selection into fragments whose lengths were randomly drawn from the distribution. Sequences shorter than 3,000 bp were left untouched.

Across all analyses, we obtained the AAI by performing protein prediction with Prodigal (version 2.6.3, parameters: “-m -p meta”) and then carrying out all-versus-all protein alignments with DIAMOND^76^ (version 2.0.15, parameter: “--sensitive”). Pairwise AAI values were then computed as the length-weighted average identity of the reciprocal best hits of pairs of scaffolds that share at least 75% of the proteins of the shortest sequence. Only matches with E-value ≤ 0.001, and query and target alignment coverage ≥ 50% were allowed.

### Marker profile database

To build a comprehensive dataset of protein profiles that could be used to identify diverse plasmids and viruses, as well as to identify provirus boundaries, we gathered protein alignments from external sources and built *de novo* clusters from a diverse collection of protein sequences. Alignments were retrieved from the following external sources: Pfam-A seed alignments (release 34.0), TIGRFAM (release 15.0), ECOD^77^ (release 20210713), EggNOG Bacteria/Archaea/Virus (version 5), VOGdb (release 206, retrieved from https://vogdb.org/), PHROG^78^, efam and efam-XC, CONJscan, double jelly-roll MCPs from Yutin, N. *et al*. (2018)^79^, *Lavidaviridae* MCPs and core proteins from Paez-Espino, D. *et al*. (2019)^80^, *Inoviridae* protein families from Roux, S. *et al*. (2019)^35^, *Leviviridae* core proteins from Callanan, J., *et al*. (2020)^70^, and RNA-dependent RNA polymerases from the RVMT dataset^51^.

*De novo* protein clusters were built from 232,031,767 protein sequences retrieved from IMG/VR v3, GTDB (release 202) species representatives, GenBank viruses (retrieved in 2021/07/06), PLSDB (release 2020_11_19), and complete metagenomic plasmids (IMG/M Taxon Object ID: 3300053491). To reduce computational overhead and prevent the formation of low diversity clusters, we first de-replicated proteins at 95% identity using MMseqs2 linclust^81^ (version 13-4511, parameters: “--kmer-per-seq 80 -c 1.0 --cluster-mode 2 --cov-mode 1 --min-seq-id 0.95”). Next, we clustered the de-replicated protein sequences with MMseqs2 cluster, requiring a minimum 80% bidirectional alignment coverage (parameters: “-s 5.5 -e 1e-5 - c 0.8 --cov-mode 0 --cluster-mode 0 --max-seqs 5000 --min-seq-id 0.5 --cluster-reassign 1”). Finally, we performed multiple sequence alignment of all the 786,782 clusters containing at least 20 proteins using Kalign^82^ (version 3.3.1). Since we wanted to maximize the profile coverage of underrepresented viral groups, we performed independent clustering of the proteins obtained from the *Nucleocytoviricota* from Schulz, F. *et al.* (2020), Asgard archaea viruses from Medvedeva, S. *et al.* (2022), archaeal tailed viruses from Liu, Y. *et al.* (2021), and unannotated domains of polyproteins from the RVMT dataset. For these datasets, we allowed clusters to contain as few as four proteins.

To identify the protein profiles that are informative for sequence classification (hereinafter markers), we measured the specificity of the 1,425,477 profiles by computing the weighted number of matches of each profile to each class (chromosome, plasmid, and virus). We first assigned weights to each sequence in such a way that the total weight of each RC within each class would be the same and that the total weight of the three classes would also be identical. Next, we converted the protein profiles into HMMs and used hmmsearch (parameter: “-E 1e-5”) to match them to Prodigal-predicted proteins from the sequence dataset. Finally, we counted the number of matches of each profile to each class taking into account the RC weights and scaled the counts within each class so that the median profile count would be the same for the three classes. Scaled counts were used to compute each profile’s Pielou’s specificity — a single summary of the profile’s specificity — and specificity measures (SPM) — which measure how specific the profiles are to each class — using the tspex^83^ tool (version 0.6.2).

To reduce the redundancy of the protein profile set, we performed a de-replication process to identify groups of profiles that target similar sets of proteins. To do that, we used the HMMs to generate artificial protein sequences with the hmmemit command (parameter: “-N 10”) and used hmmsearch (parameter: “-E 1e-5”) to align the HMMs to all artificial protein sequences. Next, to measure the empirical redundancy of all possible pairs of protein profiles, we employed the SetSimilaritySearch Python package (https://github.com/ekzhu/SetSimilaritySearch) to compute the cosine similarity of all pairs of profiles, based on the identity of their hits. Finally, we identified groups of profiles that target similar sets of proteins by clustering them using the Leiden algorithm (resolution parameter = 0.25) and selected the most specific profile of each cluster, based on their Pielou’s specificity, as its representative.

To select the markers that would be used for classification, we required protein profiles that had either Pielou’s specificity ≥ 0.4 or the maximum SPM (among the three classes) ≥ 0.75. To prevent the selection of profiles that target genes from genomic islands (which are enriched in mobile elements) as chromosome markers, we required chromosome-specific profiles to be highly prevalent by only selecting the ones that were above the median of the count distribution. For plasmid and virus markers, we required profiles to be above the first quartile of the distribution. As it was previously observed that eukaryotic sequences are prone to be misclassified as viral, we performed a negative selection of virus-specific profiles that frequently matched eukaryotic proteins. To do that, we first retrieved all the proteins that comprised eukaryotic ortholog groups in OrthoDB^84^ (version 10.1) and removed the ones that corresponded to typical viral genes, resulting in a total of 16,928,157 eukaryotic proteins. In addition, we also obtained the sequences of 159,003 proteins that were previously shown to have been horizontally transferred from viral to eukaryotic genomes^85^. We matched HMMs of virus-specific markers to the retrieved protein sequences using hmmsearch and counted the number of hits for each HMM. Profiles that matched at least 200 OrthoDB eukaryotic proteins or at least 10 horizontally transferred proteins were removed from the final marker selection. In total, 227,897 profiles were selected to be used in geNomad for distinguishing between chromosome, plasmid, and virus sequences. For benchmark purposes we repeated this process five additional times, using only the train sequences of each data split to perform the profile selection.

To assign functional annotations to the geNomad protein profiles, we used HHblits^86^ (version 3.3.0) to align them with HMMs from Pfam-A (release 35.0), TIGRFAM (release 15.0), KEGG Orthology (release 98.0), COG (release 2020), CONJscan, NCBIfam-AMRFinder (release 2022-10-11.2), and *Bacteria* and *Archaea* near-universal single-copy orthologs from BUSCO (version 5). We accepted hits with probability ≥ 90%, E-value ≤ 0.001, and target coverage ≥ 60%. Only the best hit to each database was kept, apart from Pfam, for which we allowed multiple non-overlapping hits. Names and Gene Ontology terms were assigned to geNomad markers by transferring them from the accepted Pfam, TIGRFAM, and KEGG Orthology hits. GO enrichment for each class was appraised using the Kolmogorov–Smirnov test (as implemented in the hypeR^87^ package, version 1.13.0) on lists of markers sorted by the SPM of each class. REVIGO^88^ was used to generate visualizations of the enriched GO terms.

To assign ICTV taxa to geNomad markers, we first built a protein database from viral sequences retrieved from NCBI NR (in 2022/05/19) and decorated the proteins with a custom taxdump generated from ICTV’s VMR 19 (details in Supplementary Note) using TaxonKit^89^ (version 0.11.1). We then used MMseqs2 to align geNomad’s markers to the viral protein database (parameters: “-s 8.2 -e 1e-3”) and employed taxopy (version 0.9.2, available at https://github.com/apcamargo/taxopy) to assign a single taxon to each marker by aggregating the taxonomic lineages of all the hits of each marker using the “find_majority_vote” function. Because viruses of the *Nucleocytoviricota* phylum encode homologs of bacteriophage proteins^90^, we raised the minimum fraction parameter to 0.85 to assign taxonomy to markers that were initially assigned to *Nucleocytoviricota* but matched at least one *Caudoviricetes* protein. For benchmarking purposes, we simulated taxonomic novelty by masking proteins that were similar to the ones encoded by ICTV exemplar species. This was achieved by conducting a separate round of taxonomic assignment where we ignored the hits to proteins that had ≥ 60% identity to proteins of exemplar species.

### Ecosystem distribution of markers

The distribution of geNomad’s markers across ecosystems was assessed by mapping the markers to proteins from public metagenomes and metatranscriptomes (retrieved from IMG/M on 2022-04-10) using MMseqs2 protein-profile search. The marker frequency matrix was then normalized using DESeq2^91^ (version 1.34.0), by setting the size factor of each ecosystem (according to GOLD’s ecosystem classification^92^) to the total number of proteins in it. Next, markers that mapped to less than 10 proteins were filtered out and DESeq2’s variance stabilizing transformation was employed to transform the frequency matrix. To generate the RadViz visualizations from the transformed matrix, the Radviz R library (version 0.9.3) was used.

### Classification models

To train the gene-based classifier, we first predicted the proteins encoded by the sequence fragments using prodigal-gv (version 2.7.0, parameter: “-p meta”, available at https://github.com/apcamargo/prodigal-gv) and then assigned geNomad markers to them using MMseqs2’s protein-profile search (parameters: “-s 6.4 - e 1e-3 -c 0.2 --cov-mode 1”). Next, we computed, for each sequence, a total of 25 features derived from the gene structure and marker annotation (full list and description in Supplementary Note) and used them to train a decision forest classification model with the XGBoost^93^ library (version 1.5.1, parameters: “eta=0.2, max_depth=10, n_estimators=135”). Feature selection was performed using the Boruta algorithm and SHAP importance values, as implemented in the shap-hypetune Python package (version 0.2.4, “BoostBoruta” function). Hyperparameter tuning (learning rate, tree depth, and number of trees) was performed with a random grid search (“BoostSearch” function in shap-hypetune).

The sequence-based classifier was trained using a two-step supervised contrastive learning approach^94^. In the first step we trained an IGLOO encoder to learn to produce vector representations of nucleotide sequences in such a way that sequences of the same class will tend to be clustered together and separate from sequences of different classes. To achieve this, input sequences are converted into 4-mer vectors (step size = 1) that are one-hot-encoded and zero-padded to 5,997 elements, which correspond to the number of 4-mers in a 6 kb sequence. These inputs are then fed to an IGLOO encoder, trained using the supervised contrastive loss, that produces 512-dimensional representations. In the second step, we trained a dense neural network classifier on top of the IGLOO representations using a focal loss^95^, which forces the model to focus on hard-to-classify sequences. For inference, sequences longer than 5,997 bp are split into multiple non-overlapping windows whose scores are averaged at the end of the classification. To account for class imbalance and taxonomic bias during the training of both the encoder and classifier models, sequences were weighted in accordance with their RC. For both models, training was conducted using the Adam optimizer with gradient centralization^96^. Hyperparameter tuning (k-mer size, number of IGLOO patches, number of filters in IGLOO’s convolutional layers, size of the filters, dimensionality of the classifier hidden layer) was performed with KerasTuner (version 1.1.0) using the HyperBand algorithm^97^.

The outputs of the gene-based and sequence-based classifier are aggregated by a small feedforward neural network, which uses an attention mechanism to weight the contribution of each model towards the final scores. Briefly, we trained a model that encodes in an attention matrix *A* the reliability of the gene-based classifier, estimated from the relative marker frequency within each sequence. To aggregate the results of the two classifiers, the scores generated by them are scaled according to their expected reliability encoded in *A*, and then averaged and fed to a dense layer with softmax activation.

For benchmark purposes we trained the gene-based classifier, sequence-based classifier, and the aggregator model five additional times, using the train data and the selected markers of each data split. The models used for the remaining analysis were trained with the entire dataset.

### Score calibration mechanism

To train the model underlying geNomad’s score calibration, 1,000,000 artificial communities with varying proportions of chromosome, plasmid, and virus sequences were generated by random sampling of the train dataset. For each community, scores were calibrated using an isotonic regression and the empirical composition was obtained by using geNomad to predict the most likely class of each sequence. Because isotonic regressions are dataset-specific, a regression feedforward neural network was trained to predict calibrated scores from the empirical composition and uncalibrated scores of a given community. The model was trained with the Adam optimizer and mean squared error loss.

### Provirus identification

To identify regions that correspond to proviruses within host chromosomes, geNomad employs a CRF model that was trained on a dataset of mock proviruses built from prokaryotic chromosome sequences and phage genomes. The CRF takes as input the chromosome and virus SPM values of the genes annotated with geNomad markers and computes the conditional probability of a sequence of states (chromosome or provirus). Genes are then assigned to their most likely states, forming provirus islands — that represent regions that are enriched in virus markers. To prevent having proviruses split into multiple islands due to incomplete marker coverage, provirus islands that are separated by short gene arrays (less than 6 genes or 2 chromosome markers) are merged. Next, provirus boundaries are refined by extending them to the closest tRNA (identified with ARAGORN^98^, version 1.2.41) within 5 kb and integrase (identified using MMseqs2 profile search) within 10 kb, as long as there are no chromosome markers between the original edge and the new putative coordinate. The 16 tyrosine integrase profiles used for integrase identification were manually selected from the CDD database. Finally, islands with few viral markers, which usually are not *bona fide* proviruses, are filtered out by removing the regions where the sum of the virus SPM of the markers is below a certain threshold.

### Performance benchmarks

The following tools were included in our benchmarks: geNomad (version 1.0.0), DeepMicrobeFinder (“hybrid” model, version 1.0.0), DeepVirFinder (version 1.0), PPR-Meta (version 1.1), Seeker (version 1.0.3), VIBRANT (version 1.2.1), viralVerify (version 1.1), VirSorter2 (version 2.2.3), Phigaro (version 2.3.0), PlasClass (version 0.1), and PlasX (commit 7349226). The tools were executed with default parameters and installed following the authors’ instructions, with the exception of PPR-Meta, which was executed through a Docker container. VirSorter2 was also executed with the “--include-groups dsDNAphage,NCLDV,RNA,ssDNA,lavidaviridae” parameter to measure its performance when using all classification models. To benchmark DeepMicrobeFinder we first assigned sequences to the class with the highest score, next we labeled the ones classified as “Eukaryote” or “Prokaryote” as chromosome and the ones assigned to “EukaryoteVirus” or “ProkaryoteVirus” as virus.

For the benchmarks that measured the sensitivity of virus detection across different viral and host taxa, we established cutoffs that approximated the false discovery rate of each tool to 5%. The same was done in the benchmark that measured the sensitivity of plasmid detection across different host taxa, but set the target false discovery rate to 10%, as some tools could not achieve a 5% false discovery rate regardless of the threshold. The procedure was performed to prevent overly sensitive tools (with elevated false discovery rate) from dominating the benchmarks.

### Binning of the Sand Creek Marshes metatranscriptomes

To evaluate whether geNomad is able to identify RNA virus segments that don’t encode the RdRP hallmark gene, we retrieved the raw sequencing data and the assemblies of 34 metatranscriptome samples from microbial communities from the Sand Creek Marshes (GOLD Study ID: Gs0142363)^52^ from IMG/M. Scaffolds shorter than 2 kb were discarded and the remaining sequences were classified using geNomad. Scaffolds encoding RdRP were identified by performing protein prediction with prodigal-gv and using the predicted proteins as queries to search against a database of RdRP HMM models^51^ using hmmsearch (parameter: “-E 1e-5”). Using Minimap2^99^ (version 2.24, parameters: “-N 5 -ax sr”), sequencing reads from each sample were independently mapped to sequences in a combined assembly, which was generated by concatenating the assemblies from individual samples. Then, we used samtools^100^ (version 1.16.1) to sort the mapped reads and input them into CoverM (version 0.6.1, parameter: “-m metabat”, available at https://github.com/wwood/CoverM), which measured scaffold coverage across samples. To perform an initial binning of scaffolds based on co-abundance, we employed Vamb^101^ (version 3.0.2, parameter: “-a 0.025”). Bins containing RdRP-encoding sequences were refined by retaining only the scaffolds that presented high correlation to the coverage of the RdRP scaffold (Pearson correlation coefficient ≥ 0.95). To prevent spurious correlations, we only considered RdRP-encoding sequences with high prevalence (coverage > 0 in at least 20% of the samples).

### Metagenomic survey and phylogenetic analysis of giant viruses

From a set of 28,865 metagenomes (retrieved on 2022-04-10 from IMG/M) we selected scaffolds longer than 50 kb that were classified as viruses by geNomad and subjected them to further processing using GVClass (available at https://github.com/NeLLi-team/gvclass/), a framework that identifies giant viruses and assigns them to taxonomic lineages using a phylogenetic placement approach. Briefly, we identified nine conserved giant virus orthologous groups (GVOGs)^102^ using hmmsearch and used these GVOGs as queries for DIAMOND searches against databases of the respective GVOGs, which were built from a representative set of bacteria, archaea, eukaryotes, and viruses. We extracted the top 100 hits, combined them with the query sequences, and aligned them with MAFFT^103^ (version 7.490). The alignments were then trimmed with trimAl^104^ (version 1.4, parameter: “-gt 0.1”) and used to build a phylogenetic tree with FastTree^105^ (version 2.1.11, parameters: “-spr 4 -mlacc 2 -slownni -lg”). To determine the final classification, we identified the nearest neighbor in the tree using branch lengths and the existing taxonomic string for that reference genome. The taxonomic strings from all identified nearest neighbors were then compared at different taxonomic levels (genus, family, order, class, phylum) to yield the final classification at the lowest taxonomic level on which all nearest neighbors were in agreement.

To measure the phylogenetic diversity (PD) gained by identifying giant viruses with geNomad, we extracted DNA PolB orthologs encoded by these sequences and by genomes from two external sources (Schulz et al., 2020, Aylward et al., 2021; only sequences on scaffolds longer than 50 kb were considered) using the DNA PolB HMM model from the GVOG database (GVOGm0054). These protein sequences were aligned with MAFFT, trimmed with trimAl, and used to build a phylogenetic tree with FastTree. We then performed separate alignments and built trees for each of the orders in the *Nucleocytoviricota*, with and without geNomad contigs. The increase in PD was then determined as the fold difference between the sum of the branch lengths for each viral order after adding the giant viruses identified with geNomad.

To build the phylogenetic tree that included the giant viruses identified from soil metagenomes using geNomad, we employed a representative set of giant viruses from Aylward et al. (2021) and added additional GVMAGs recovered from soil samples in Schulz et al. (2020). The sequences of the seven predominantly vertically inherited GVOGs were identified across all scaffolds using hmmsearch, aligned using MAFFT, and trimmed with trimA. Subsequently, a concatenated alignment was used as input to reconstruct a phylogenetic tree with IQ-TREE^106^ (version 2.2.0.3, parameters: “-m LG+F+I+G4”).

## Code and data availability

geNomad is an open source software and its code can be found at https://github.com/apcamargo/genomad. Multiple sequence alignments, HMMs, and a MMseqs2 database of geNomad’s markers are available at https://doi.org/10.5281/zenodo.7586412. The code used to build the taxonomically annotated viral protein database can be found at https://github.com/apcamargo/ictv-mmseqs2-protein-database and the database can be downloaded from https://doi.org/10.5281/zenodo.6574913. Reference sequences utilized for training and evaluation, the list of *P. aeruginosa* genomes used to build the pangenome, giant virus sequences discovered in metagenomes, and the code applied to train geNomad’s neural network and conditional random field models can be downloaded from https://doi.org/10.5281/zenodo.7697490.

## Acknowledgements

We thank Colin Hill, Lee Call, Mart Krupovic, Uri Neri, Eduardo P. C. Rocha, and Ahmed A. Zayed for providing genomic and multiple sequence alignment data that were used to assemble geNomad’s marker dataset. We extend our appreciation to Adrielle A. Vasconcelos for her assistance in establishing geNomad’s name and visual identity.The JGI Award DOIs of the metagenomes where giant viruses were identified are available at https://doi.org/10.5281/zenodo.7697490. The work conducted by the U.S. Department of Energy Joint Genome Institute (https://ror.org/04xm1d337), and the National Energy Research Scientific Computing Center (NERSC) (https://ror.org/05v3mvq14), which are DOE Office of Science User Facilities operated under Contract No. DE-AC02-05CH11231. This work also received support from the Genomic Science Program in the U.S. Department of Energy, Office of Science, Office of Biological and Environmental Research (BER) (89233218CNA000001 to L.A.N.L., DE-AC05-00OR22725 to O.R.N.L., DE-AC05-76RL01830 to P.N.N.L.). We also used computational resources from the Exascale Computing Project (17-SC-20-SC), a collaborative effort of the U.S. Department of Energy Office of Science and the National Nuclear Security Administration.

## References

1. Rodríguez-Beltrán, J., DelaFuente, J., León-Sampedro, R., MacLean, R. C. & San Millán, Á. Beyond horizontal gene transfer: the role of plasmids in bacterial evolution. Nat. Rev. Microbiol. 19, 347–359 (2021).

2. Suttle, C. A. Viruses in the sea. Nature 437, 356–361 (2005).

3. Ochman, H., Lawrence, J. G. & Groisman, E. A. Lateral gene transfer and the nature of bacterial innovation. Nature 405, 299–304 (2000).

4. de la Cruz, F. & Davies, J. Horizontal gene transfer and the origin of species: lessons from bacteria. Trends Microbiol. 8, 128–133 (2000).

5. Smalla, K., Jechalke, S. & Top, E. M. Plasmid Detection, Characterization, and Ecology. Microbiol. Spectr. 3, 3.1.17 (2015).

6. Yu, M. K., Fogarty, E. C. & Eren, A. M. The genetic and ecological landscape of plasmids in the human gut. bioRxiv (2020) doi:10.1101/2020.11.01.361691.

7. Eraslan, G., Avsec, Ž., Gagneur, J. & Theis, F. J. Deep learning: new computational modelling techniques for genomics. Nat. Rev. Genet. 20, 389–403 (2019).

8. Ren, J. et al. Likelihood Ratios for Out-of-Distribution Detection. in Proceedings of the 33rd International Conference on Neural Information Processing Systems (Curran Associates Inc., 2019).

9. Fouts, D. E. Phage_Finder: Automated identification and classification of prophage regions in complete bacterial genome sequences. Nucleic Acids Res. 34, 5839–5851 (2006).

10. Camargo, A. P. et al. IMG/VR v4: an expanded database of uncultivated virus genomes within a framework of extensive functional, taxonomic, and ecological metadata. Nucleic Acids Res. 51, D733–D743 (2023).

11. Sourkov, V. IGLOO: Slicing the Features Space to Represent Sequences. arXiv (2018) doi:10.48550/ARXIV.1807.03402.

12. Camargo, A. P., Sourkov, V., Pereira, G. A. G. & Carazzolle, M. F. RNAsamba: neural network-based assessment of the protein-coding potential of RNA sequences. NAR Genomics Bioinforma. 2, lqz024 (2020).

13. Hyatt, D. et al. Prodigal: prokaryotic gene recognition and translation initiation site identification. BMC Bioinformatics 11, 119 (2010).

14. Yutin, N. et al. Analysis of metagenome-assembled viral genomes from the human gut reveals diverse putative CrAss-like phages with unique genomic features. Nat. Commun. 12, 1044 (2021).

15. Schulz, F. et al. Giant virus diversity and host interactions through global metagenomics. Nature 578, 432–436 (2020).

16. Steinegger, M. & Söding, J. MMseqs2 enables sensitive protein sequence searching for the analysis of massive data sets. Nat. Biotechnol. 35, 1026–1028 (2017).

17. Walker, P. J. et al. Recent changes to virus taxonomy ratified by the International Committee on Taxonomy of Viruses (2022). Arch. Virol. 167, 2429–2440 (2022).

18. Zayed, A. A. et al. efam: an *e*xpanded, metaproteome-supported HMM profile database of viral protein *fam*ilies. Bioinformatics 37, 4202–4208 (2021).

19. Huerta-Cepas, J. et al. eggNOG 5.0: a hierarchical, functionally and phylogenetically annotated orthology resource based on 5090 organisms and 2502 viruses. Nucleic Acids Res. 47, D309–D314 (2019).

20. Mistry, J. et al. Pfam: The protein families database in 2021. Nucleic Acids Res. 49, D412–D419 (2021).

21. Haft, D. H., Selengut, J. D. & White, O. The TIGRFAMs database of protein families. Nucleic Acids Res. 31, 371–373 (2003).

22. Kanehisa, M. & Goto, S. KEGG: Kyoto Encyclopedia of Genes and Genomes. Nucleic Acids Res. 28, 27–30 (2000).

23. Galperin, M. Y. et al. COG database update: focus on microbial diversity, model organisms, and widespread pathogens. Nucleic Acids Res. 49, D274–D281 (2021).

24. Cury, J., Abby, S. S., Doppelt-Azeroual, O., Néron, B. & Rocha, E. P. C. Identifying Conjugative Plasmids and Integrative Conjugative Elements with CONJscan. in Horizontal Gene Transfer: Methods and Protocols (ed. de la Cruz, F.) 265–283 (Springer US, 2020). doi:10.1007/978-1-4939-9877-7_19.

25. Feldgarden, M. et al. AMRFinderPlus and the Reference Gene Catalog facilitate examination of the genomic links among antimicrobial resistance, stress response, and virulence. Sci. Rep. 11, 12728 (2021).

26. Manni, M., Berkeley, M. R., Seppey, M., Simão, F. A. & Zdobnov, E. M. BUSCO Update: Novel and Streamlined Workflows along with Broader and Deeper Phylogenetic Coverage for Scoring of Eukaryotic, Prokaryotic, and Viral Genomes. Mol. Biol. Evol. 38, 4647–4654 (2021).

27. Hou, S., Cheng, S., Chen, T., Fuhrman, J. A. & Sun, F. DeepMicrobeFinder sorts metagenomes into prokaryotes, eukaryotes and viruses, with marine applications. bioRxiv (2021) doi:10.1101/2021.10.26.466018.

28. Fang, Z. et al. PPR-Meta: a tool for identifying phages and plasmids from metagenomic fragments using deep learning. GigaScience 8, giz066 (2019).

29. Pellow, D., Mizrahi, I. & Shamir, R. PlasClass improves plasmid sequence classification. PLOS Comput. Biol. 16, e1007781 (2020).

30. Antipov, D., Raiko, M., Lapidus, A. & Pevzner, P. A. METAVIRALSPADES: assembly of viruses from metagenomic data. Bioinformatics 36, 4126–4129 (2020).

31. Guo, J. et al. VirSorter2: a multi-classifier, expert-guided approach to detect diverse DNA and RNA viruses. Microbiome 9, 37 (2021).

32. Kieft, K., Zhou, Z. & Anantharaman, K. VIBRANT: automated recovery, annotation and curation of microbial viruses, and evaluation of viral community function from genomic sequences. Microbiome 8, 90 (2020).

33. Auslander, N., Gussow, A. B., Benler, S., Wolf, Y. I. & Koonin, E. V. Seeker: alignment-free identification of bacteriophage genomes by deep learning. Nucleic Acids Res. 48, e121–e121 (2020).

34. Ren, J. et al. Identifying viruses from metagenomic data using deep learning. Quant. Biol. 8, 64–77 (2020).

35. Roux, S. et al. Cryptic inoviruses revealed as pervasive in bacteria and archaea across Earth’s biomes. Nat. Microbiol. 4, 1895–1906 (2019).

36. Wagner, P. L. & Waldor, M. K. Bacteriophage control of bacterial virulence. Infect. Immun. 70, 3985– 3993 (2002).

37. Bondy-Denomy, J. et al. Prophages mediate defense against phage infection through diverse mechanisms. ISME J. 10, 2854–2866 (2016).

38. Carey, J. N. et al. Phage integration alters the respiratory strategy of its host. eLife 8, e49081 (2019).

39. Zhou, Y., Liang, Y., Lynch, K. H., Dennis, J. J. & Wishart, D. S. PHAST: A Fast Phage Search Tool. Nucleic Acids Res. 39, W347–W352 (2011).

40. Mageeney, C. M. et al. New candidates for regulated gene integrity revealed through precise mapping of integrative genetic elements. Nucleic Acids Res. 48, 4052–4065 (2020).

41. Zünd, M. et al. High throughput sequencing provides exact genomic locations of inducible prophages and accurate phage-to-host ratios in gut microbial strains. Microbiome 9, 77 (2021).

42. Rousset, F. et al. Phages and their satellites encode hotspots of antiviral systems. Cell Host Microbe 30, 740–753.e5 (2022).

43. Starikova, E. V. et al. Phigaro: high-throughput prophage sequence annotation. Bioinformatics 36, 3882– 3884 (2020).

44. Nayfach, S. et al. CheckV assesses the quality and completeness of metagenome-assembled viral genomes. Nat. Biotechnol. 39, 578–585 (2021).

45. Gautreau, G. et al. PPanGGOLiN: Depicting microbial diversity via a partitioned pangenome graph. PLOS Comput. Biol. 16, e1007732 (2020).

46. LeRoux, M. et al. The DarTG toxin-antitoxin system provides phage defence by ADP-ribosylating viral DNA. Nat. Microbiol. 7, 1028–1040 (2022).

47. Doron, S. et al. Systematic discovery of antiphage defense systems in the microbial pangenome. Science 359, eaar4120 (2018).

48. Tesson, F. et al. Systematic and quantitative view of the antiviral arsenal of prokaryotes. Nat. Commun. 13, 2561 (2022).

49. Edgar, R. C. et al. Petabase-scale sequence alignment catalyses viral discovery. Nature 602, 142–147 (2022).

50. Zayed, A. A. et al. Cryptic and abundant marine viruses at the evolutionary origins of Earth’s RNA virome. Science 376, 156–162 (2022).

51. Neri, U. et al. Expansion of the global RNA virome reveals diverse clades of bacteriophages. Cell 185, 4023–4037.e18 (2022).

52. H. Vineis, J. Nutrient influence on microbial structure and function within salt marsh sediments. (Northeastern University, 2022).

53. Koonin, E. V. & Yutin, N. Origin and Evolution of Eukaryotic Large Nucleo-Cytoplasmic DNA Viruses. Intervirology 53, 284–292 (2010).

54. Schulz, F. et al. Giant viruses with an expanded complement of translation system components. Science 356, 82–85 (2017).

55. Bäckström, D. et al. Virus Genomes from Deep Sea Sediments Expand the Ocean Megavirome and Support Independent Origins of Viral Gigantism. mBio 10, e02497–18 (2019).

56. Chen, I.-M. A. et al. The IMG/M data management and analysis system v.7: content updates and new features. Nucleic Acids Res. 51, D723–D732 (2023).

57. Gallot-Lavallée, L., Blanc, G. & Claverie, J.-M. Comparative Genomics of Chrysochromulina Ericina Virus and Other Microalga-Infecting Large DNA Viruses Highlights Their Intricate Evolutionary Relationship with the Established Mimiviridae Family. J. Virol. 91, e00230–17 (2017).

58. Schulz, F. et al. Hidden diversity of soil giant viruses. Nat. Commun. 9, 4881 (2018).

59. Lang, A. S., Zhaxybayeva, O. & Beatty, J. T. Gene transfer agents: phage-like elements of genetic exchange. Nat. Rev. Microbiol. 10, 472–482 (2012).

60. Ghequire, M. G. K. & Mot, R. D. The Tailocin Tale: Peeling off Phage Tails. Trends Microbiol. 23, 587–590 (2015).

61. Johansen, J. et al. Genome binning of viral entities from bulk metagenomics data. Nat. Commun. 13, 965 (2022).

62. Schulz, F. et al. Advantages and Limits of Metagenomic Assembly and Binning of a Giant Virus. mSystems 5, e00048–20 (2020).

63. Cury, J., Touchon, M. & Rocha, E. P. C. Integrative and conjugative elements and their hosts: composition, distribution and organization. Nucleic Acids Res. 45, 8943–8956 (2017).

64. Parks, D. H. et al. GTDB: an ongoing census of bacterial and archaeal diversity through a phylogenetically consistent, rank normalized and complete genome-based taxonomy. Nucleic Acids Res. 50, D785–D794 (2022).

65. Schmartz, G. P. et al. PLSDB: advancing a comprehensive database of bacterial plasmids. Nucleic Acids Res. 50, D273–D278 (2022).

66. Alexander, H. et al. Eukaryotic genomes from a global metagenomic dataset illuminate trophic modes and biogeography of ocean plankton. bioRxiv (2021) doi:10.1101/2021.07.25.453713.

67. Traag, V. A., Waltman, L. & van Eck, N. J. From Louvain to Leiden: guaranteeing well-connected communities. Sci. Rep. 9, 5233 (2019).

68. Eddy, S. R. Accelerated Profile HMM Searches. PLOS Comput. Biol. 7, e1002195 (2011).

69. Roux, S. et al. IMG/VR v3: an integrated ecological and evolutionary framework for interrogating genomes of uncultivated viruses. Nucleic Acids Res. 49, D764–D775 (2021).

70. Callanan, J. et al. Expansion of known ssRNA phage genomes: From tens to over a thousand. Sci. Adv. 6, eaay5981 (2020).

71. Medvedeva, S. et al. Three families of Asgard archaeal viruses identified in metagenome-assembled genomes. Nat. Microbiol. 7, 962–973 (2022).

72. Liu, Y. et al. Diversity, taxonomy, and evolution of archaeal viruses of the class Caudoviricetes. PLOS Biol. 19, e3001442 (2021).

73. Roux, S. et al. Minimum Information about an Uncultivated Virus Genome (MIUViG). Nat. Biotechnol. 37, 29–37 (2019).

74. Zhang, Z., Schwartz, S., Wagner, L. & Miller, W. A Greedy Algorithm for Aligning DNA Sequences. J. Comput. Biol. 7, 203–214 (2000).

75. Parks, D. H. et al. A complete domain-to-species taxonomy for Bacteria and Archaea. Nat. Biotechnol. 38, 1079–1086 (2020).

76. Buchfink, B., Reuter, K. & Drost, H.-G. Sensitive protein alignments at tree-of-life scale using DIAMOND. Nat. Methods 18, 366–368 (2021).

77. Cheng, H. et al. ECOD: An Evolutionary Classification of Protein Domains. PLoS Comput. Biol. 10, e1003926 (2014).

78. Terzian, P., et al. PHROG: families of prokaryotic virus proteins clustered using remote homology. NAR Genomics Bioinforma. 3, lqab067 (2021).

79. Yutin, N., Bäckström, D., Ettema, T. J. G., Krupovic, M. & Koonin, E. V. Vast diversity of prokaryotic virus genomes encoding double jelly-roll major capsid proteins uncovered by genomic and metagenomic sequence analysis. Virol. J. 15, 67 (2018).

80. Paez-Espino, D. et al. Diversity, evolution, and classification of virophages uncovered through global metagenomics. Microbiome 7, 157 (2019).

81. Steinegger, M. & Söding, J. Clustering huge protein sequence sets in linear time. Nat. Commun. 9, 2542 (2018).

82. Lassmann, T. Kalign 3: multiple sequence alignment of large datasets. Bioinformatics 36, 1928–1929 (2020).

83. Camargo, A. P., Vasconcelos, A. A., Fiamenghi, M. B., Pereira, G. A. G. & Carazzolle, M. F. tspex: a tissue-specificity calculator for gene expression data. Res. Sq. (2020) doi:10.21203/rs.3.rs-51998/v1.

84. Kriventseva, E. V. et al. OrthoDB v10: sampling the diversity of animal, plant, fungal, protist, bacterial and viral genomes for evolutionary and functional annotations of orthologs. Nucleic Acids Res. 47, D807–D811 (2019).

85. Irwin, N. A. T., Pittis, A. A., Richards, T. A. & Keeling, P. J. Systematic evaluation of horizontal gene transfer between eukaryotes and viruses. Nat. Microbiol. 7, 327–336 (2022).

86. Remmert, M., Biegert, A., Hauser, A. & Söding, J. HHblits: lightning-fast iterative protein sequence searching by HMM-HMM alignment. Nat. Methods 9, 173–175 (2012).

87. Federico, A. & Monti, S. hypeR: an R package for geneset enrichment workflows. Bioinformatics 36, 1307–1308 (2020).

88. Supek, F., Bošnjak, M., Škunca, N. & Šmuc, T. REVIGO Summarizes and Visualizes Long Lists of Gene Ontology Terms. PLOS ONE 6, e21800 (2011).

89. Shen, W. & Ren, H. TaxonKit: A practical and efficient NCBI taxonomy toolkit. J. Genet. Genomics 48, 844–850 (2021).

90. Mönttinen, H. A. M., Bicep, C., Williams, T. A. & Hirt, R. P. The genomes of nucleocytoplasmic large DNA viruses: viral evolution writ large. *Microb*. Genomics 7, 000649 (2021).

91. Love, M. I., Huber, W. & Anders, S. Moderated estimation of fold change and dispersion for RNA-seq data with DESeq2. Genome Biol. 15, 550 (2014).

92. Mukherjee, S. et al. Twenty-five years of Genomes OnLine Database (GOLD): data updates and new features in v.9. Nucleic Acids Res. 51, D957–D963 (2023).

93. Chen, T. & Guestrin, C. XGBoost: A Scalable Tree Boosting System. in Proceedings of the 22nd ACM SIGKDD International Conference on Knowledge Discovery and Data Mining 785–794 (ACM, 2016). doi:10.1145/2939672.2939785.

94. Khosla, P. et al. Supervised Contrastive Learning. in Advances in Neural Information Processing Systems (eds. Larochelle, H., Ranzato, M., Hadsell, R., Balcan, M. F. & Lin, H.) vol. 33 18661–18673 (Curran Associates, Inc., 2020).

95. Lin, T.-Y., Goyal, P., Girshick, R., He, K. & Dollár, P. Focal Loss for Dense Object Detection. IEEE Trans. Pattern Anal. Mach. Intell. 42, 318–327 (2020).

96. Yong, H., Huang, J., Hua, X. & Zhang, L. Gradient Centralization: A New Optimization Technique for Deep Neural Networks. in Computer Vision – ECCV 2020 (eds. Vedaldi, A., Bischof, H., Brox, T. & Frahm, J.-M.) vol. 12346 635–652 (Springer International Publishing, 2020).

97. Li, L., Jamieson, K., DeSalvo, G., Rostamizadeh, A. & Talwalkar, A. Hyperband: A Novel Bandit-Based Approach to Hyperparameter Optimization. J Mach Learn Res 18, 6765–6816 (2017).

98. Laslett, D. & Canback, B. ARAGORN, a program to detect tRNA genes and tmRNA genes in nucleotide sequences. Nucleic Acids Res. 32, 11–16 (2004).

99. Li, H. Minimap2: pairwise alignment for nucleotide sequences. Bioinformatics 34, 3094–3100 (2018).

100. Li, H. et al. The Sequence Alignment/Map format and SAMtools. Bioinformatics 25, 2078–2079 (2009).

101. Nissen, J. N. et al. Improved metagenome binning and assembly using deep variational autoencoders. Nat. Biotechnol. 39, 555–560 (2021).

102. Aylward, F. O., Moniruzzaman, M., Ha, A. D. & Koonin, E. V. A phylogenomic framework for charting the diversity and evolution of giant viruses. PLOS Biol. 19, e3001430 (2021).

103. Katoh, K. & Standley, D. M. MAFFT Multiple Sequence Alignment Software Version 7: Improvements in Performance and Usability. Mol. Biol. Evol. 30, 772–780 (2013).

104. Capella-Gutiérrez, S., Silla-Martínez, J. M. & Gabaldón, T. trimAl: a tool for automated alignment trimming in large-scale phylogenetic analyses. Bioinformatics 25, 1972–1973 (2009).

105. Price, M. N., Dehal, P. S. & Arkin, A. P. FastTree 2 – Approximately Maximum-Likelihood Trees for Large Alignments. PLOS ONE 5, e9490 (2010).

106. Minh, B. Q. et al. IQ-TREE 2: New Models and Efficient Methods for Phylogenetic Inference in the Genomic Era. Mol. Biol. Evol. 37, 1530–1534 (2020).

