## Supplementary Note for "You can move, but you can’t hide: identification of mobile genetic elements with geNomad"

### Marker-based classification features

To classify a given sequence into chromosome, plasmid, or virus with the marker-based classifier, geNomad performs gene prediction using prodigal-gv and annotates the predicted proteins by aligning them to geNomad's markers using MMseqs2. From the sequence's gene structure, RBS motifs, and the identity of the markers that were assigned to its proteins, a total of 25 informative features are computed and used as input for the classification model. Below we list and describe each one of these features:

- **`strand_switch_rate`:** The fraction of genes located on a different strand from the gene upstream.
- **`coding_density`:** Sum of the lengths of all the protein-coding regions (in base pairs) divided by the total sequence length.
- **`no_rbs_freq`:** Fraction of genes without a detectable RBS motif.
- **`sd_bacteroidetes_rbs_freq`:** Fraction of genes predicted to have a Bacteroidetes Shine-Dalgarno RBS motif (TAA, TAAA, TAAAA, TAAAT, TAAAAA, TAAAAT).
- **`sd_canonical_rbs_freq`:** Fraction of genes predicted to have a canonical Shine-Dalgarno RBS motif (3Base/5BMM, 4Base/6BMM, AGG, AGGA, AGGA/GGAG/GAGG, AGGAG, AGGAG/GGAGG, AGGAG(G)/GGAGG, AGGAGG, AGxAG, AGxAGG/AGGxGG, GAG, GAGG, GAGGA, GGA, GGA/GAG/AGG, GGAG, GGAG/GAGG, GGAGG, GGAGGA, GGxGG).
- **`tatata_rbs_freq`:** Fraction of genes predicted to have a TATATA RBS motif (ATA, ATAT, ATATA, ATATAT, TAT, TATA, TATAT, TATATA).
- **`cc_marker_freq`:** Number of genes assigned to the CC specificity class (high chromosome SPM, low plasmid SPM, low virus SPM) divided by the total number of genes.
- **`cp_marker_freq`:** Number of genes assigned to the CP specificity class (high chromosome SPM, medium plasmid SPM, low virus SPM) divided by the total number of genes.
- **`cv_marker_freq`:** Number of genes assigned to the CV specificity class (high chromosome SPM, low plasmid SPM, medium virus SPM) divided by the total number of genes.
- **`pc_marker_freq`:** Number of genes assigned to the PC specificity class (medium chromosome SPM, high plasmid SPM, low virus SPM) divided by the total number of genes.
- **`pp_marker_freq`:** Number of genes assigned to the PP specificity class (low chromosome SPM, high plasmid SPM, low virus SPM) divided by the total number of genes.
- **`pv_marker_freq`:** Number of genes assigned to the PV specificity class (low chromosome SPM, high plasmid SPM, medium virus SPM) divided by the total number of genes.
- **`vc_marker_freq`:** Number of genes assigned to the VC specificity class (medium chromosome SPM, low plasmid SPM, high virus SPM) divided by the total number of genes.
- **`vp_marker_freq`:** Number of genes assigned to the VP specificity class (low chromosome SPM, medium plasmid SPM, high virus SPM) divided by the total number of genes.

- **vv_marker_freq:** Number of genes assigned to the VV specificity class (low chromosome SPM, low plasmid SPM, high virus SPM) divided by the total number of genes.
- **c_marker_freq:** Total chromosome marker frequency (CC + CP + CV).
- **p_marker_freq:** Total plasmid marker frequency (PC + PP + PV).
- **v_marker_freq:** Total virus marker frequency (VC + VP + VV).
- **median_c_spm:** Median chromosome SPM across all annotated genes.
- **median_p_spm:** Median plasmid SPM across all annotated genes.
- **median_v_spm:** Median virus SPM across all annotated genes.
- **v_vs_c_score_logistic:** A sigmoid function is applied to a compound score($\sum_{i=1}^{n} V\,SPM_i - C\,SPM_i$) to put it in the $[0-1]$ range.
- **v_vs_p_score_logistic:** A sigmoid function is applied to a compound score($\sum_{i=1}^{n} V\,SPM_i - P\,SPM_i$) to put it in the $[0-1]$ range.
- **p_vs_v_score_logistic:** A sigmoid function is applied to a compound score($\sum_{i=1}^{n} P\,SPM_i - V\,SPM_i$) to put it in the $[0-1]$ range.
- **gv_marker_freq:** Number of genes annotated with giant virus markers divided by the total number of genes.

Notes:

1. Predicted RBS motifs are extracted from prodigal-gv's gene prediction.
2. Each profile has three associated SPM values that range from 0 to 1 and measure how specific that profile is to each one of the three classes (chromosome, plasmid, and virus).
3. Markers were assigned to the nine specificity classes (CC, CP, CV, PC, PP, PV, VC, VP, and VV) based on their SPM values. Briefly, we used the "binned_statistic_dd" function from the SciPy Python library (version 1.7.3) to divide the three-dimensional SPM space into 125 equally sized bins. Next, each marker was assigned to a bin based on its SPM profile, so that all the markers within a given bin had similar chromosome, plasmid, and virus SPMs. Finally, we manually labeled each bin, and the markers within it, with the nine specificity classes, depending on their SPM profiles.
4. To label profiles as giant virus markers, we treated giant viruses (*Nucleocytoviricota*, *Pandoravirus*, *Mollivirus*, *Pithoviridae*, *Naldaviricetes*) as a fourth class, separate from all other viruses, and recomputed SPM values. Profiles with giant virus SPM ≥ 0.94 were considered giant virus markers. This threshold was picked based on the SPM of profiles of known *Megaviricetes* capsid proteins.
