## Supplementary Figures Legends for "You can move, but you can’t hide: identification of mobile genetic elements with geNomad"

**Supplementary Figure 1 | Global sequence representations generated by the IGLOO encoder are used for sequence classification. (A)** The IGLOO encoder applies 128 independent convolutions to the one-hot-encoded sequence to create a feature map, from which four random slices are taken and concatenated to generate patches that encode long-distance relationships within the sequence. **(B)** A total of 2,100 patches are used to weight different parts of the feature map in a transformer-like self-attention mechanism that results in a high-dimensional sequence representation. The encoder was trained using a supervised contrastive loss function, which optimizes the separation of the three classes (chromosome, plasmid, and virus) in the embedding space. **(C)** To classify sequences, the sequence representations generated by the IGLOO encoder are fed to a dense neural network trained with focal loss to account for class imbalance.

**Supplementary Figure 2 | Sample composition can be leveraged to calibrate classification scores to approximate probabilities. (A)** The false positive rates of a set of classifications depend on the sample's underlying composition. In typical metagenomes, where cellular sequences outnumber viral sequences, the fraction of false positives within scaffolds classified as viral is higher than in a virome. **(B)** The mean absolute error (MAE) of the score calibration model (*y*-axis) is highly dependent on the number of sequences in the sample (*x*-axis), as larger samples will result in more accurate estimates of the underlying sample composition. **(C)** The calibration model tends to increase the scores of a given class when it is abundant in the sample and reduce the scores when the class is rare. **(D)** The relative frequency of a given class in the sample (*x*-axis) contributes positively to the model output (*y*-axis, quantified using SHAP) when that class is abundant in the sample and negatively when the class is rare. **(E)** The pre-calibration score of a given class in the sample (*x*-axis) contributes positively to the model output (*y*-axis, quantified using SHAP) when the initial score is high and negatively when the initial score is low.

**Supplementary Figure 3 | Assigning viral taxa using geNomad's markers. (A)** To assign viral sequences to specific taxa, geNomad utilizes a best-hit approach to initially assign the genes encoded by these sequences to markers. **(B)** Each gene is subsequently classified based on the taxonomic lineage of the assigned marker. Different genes within the sequence might be assigned to different lineages. **(C)** To establish a single sequence-level taxonomy, geNomad aggregates the lineages of all the markers using a weighted majority vote approach. This approach determines the support for each taxon at each taxonomic rank by summing the bitscores of all genes assigned to that taxon. The sequence is then assigned to the most specific taxon that is supported by at least 50% of the total bitscore of the sequence.

**Supplementary Figure 4 | geNomad's marker dataset was built by gathering dereplicated protein profiles from several sources and measuring their specificity to chromosomes, plasmids, and viruses. (A)** Number of protein profile clusters obtained by varying the clustering granularity (Leiden's resolution parameter). The value chosen for dereplication (0.25) is indicated in blue. **(B)** UpSet plot showing the overlap of different protein profile datasets in the dereplication process. The overlap between a given pair of datasets was measured as the number of protein profile clusters that contained profiles from both. **(C)** Ternary plot showing the specificity of protein profiles (circles) prior to dereplication (n = 470,039). Colors represent the marker density in a region of the plot.

**Supplementary Figure 5 | geNomad can detect plasmids and viruses with low identity to the training data even if they encode few or no markers. (A)** Length distributions of the sequence fragments used to train geNomad and to evaluate classification performance of multiple tools. Sequence length (*x*-axis) is represented in log scale. **(B)** geNomad's classification performance on plasmids (left) and viruses (right) with varying degrees of similarity to sequences in the train data (bins in the *x*-axis). Similarity to the train data was assessed by computing average amino acid identities to the sequences in the train data. **(C)** geNomad's classification performance on plasmids (left) and viruses (right) with varying marker frequency (fraction of genes assigned to a geNomad marker). For each interval, performance was measured across five pairs of train/test sets (leave-one-group-out strategy). **(D)** Score calibration improves classification performance for both plasmids and viruses across all length ranges. Classification performance was measured using the Matthews correlation coefficient (MCC).

**Supplementary Figure 6 | geNomad outperforms other tools in identifying proviruses in the *Pseudomonas aeruginosa* pangenome. (A)** Distribution of the contamination estimates of multiple provirus-identification tools, measured at the gene-level for each provirus. Contamination was measured as the number of core genes, as determined by PPanGGOLiN, in the provirus. The number of detected provirus and the median contamination of each tool are displayed below the graph. **(B)** Defense system-encoding proviral regions demarcated with multiple tools in *P. aeruginosa* genomes. Shell and cloud genes are shown in light grey and core genes (putative contamination) are shown in dark gray. Genes that are part of defense systems are in orange. Integrase genes are in blue. tRNA loci are indicated by red arrows. GenBank accessions are shown within parenthesis. Phigaro did not detect any provirus within the 2,370,782–2,449,616 bp region in the NZ_CP078009.1 sequence.
