## Supplementary figures and images for "You can move, but you can’t hide: identification of mobile genetic elements with geNomad"

### Supplementary Figure 1

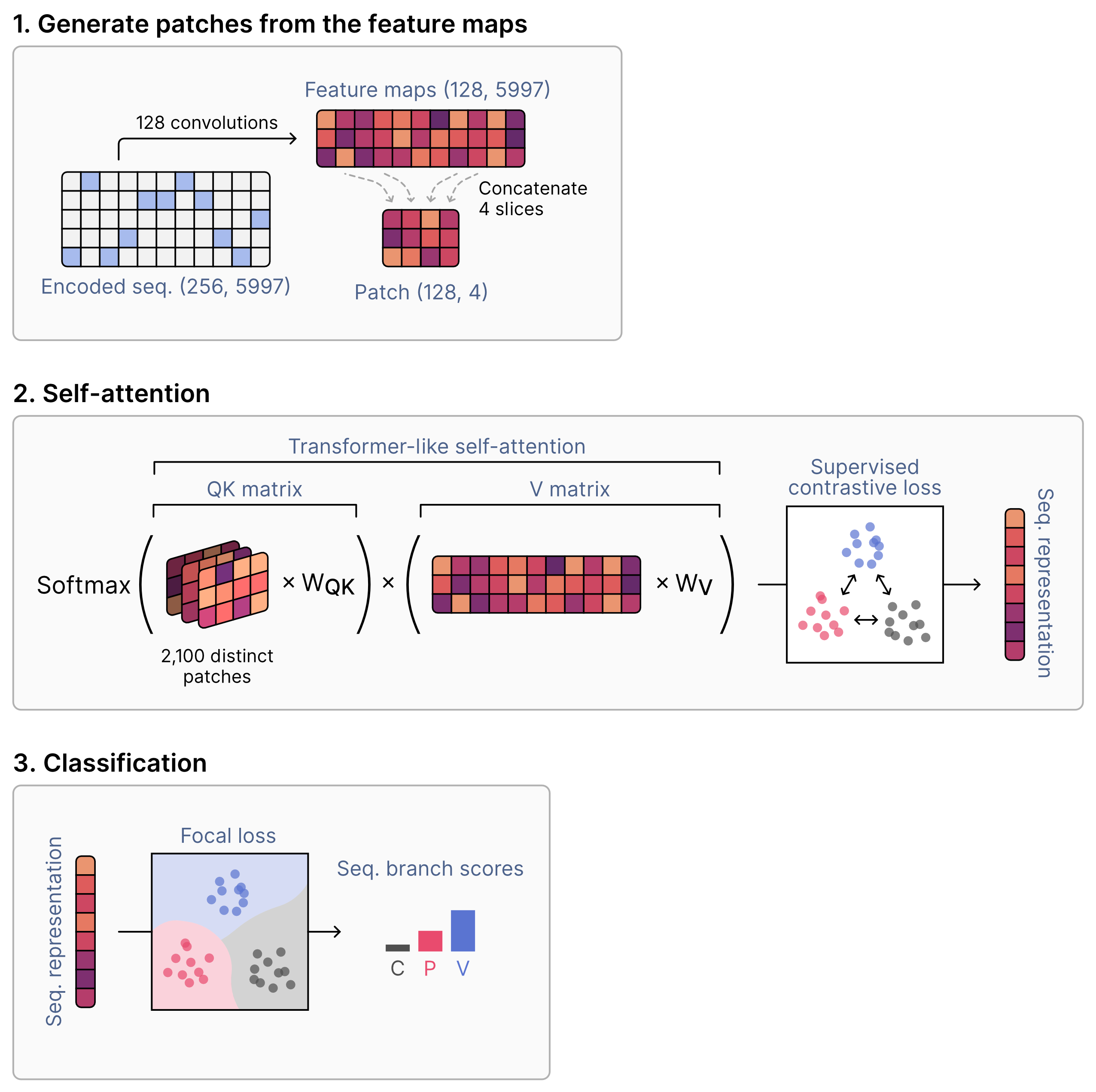

### Supplementary Figure 2

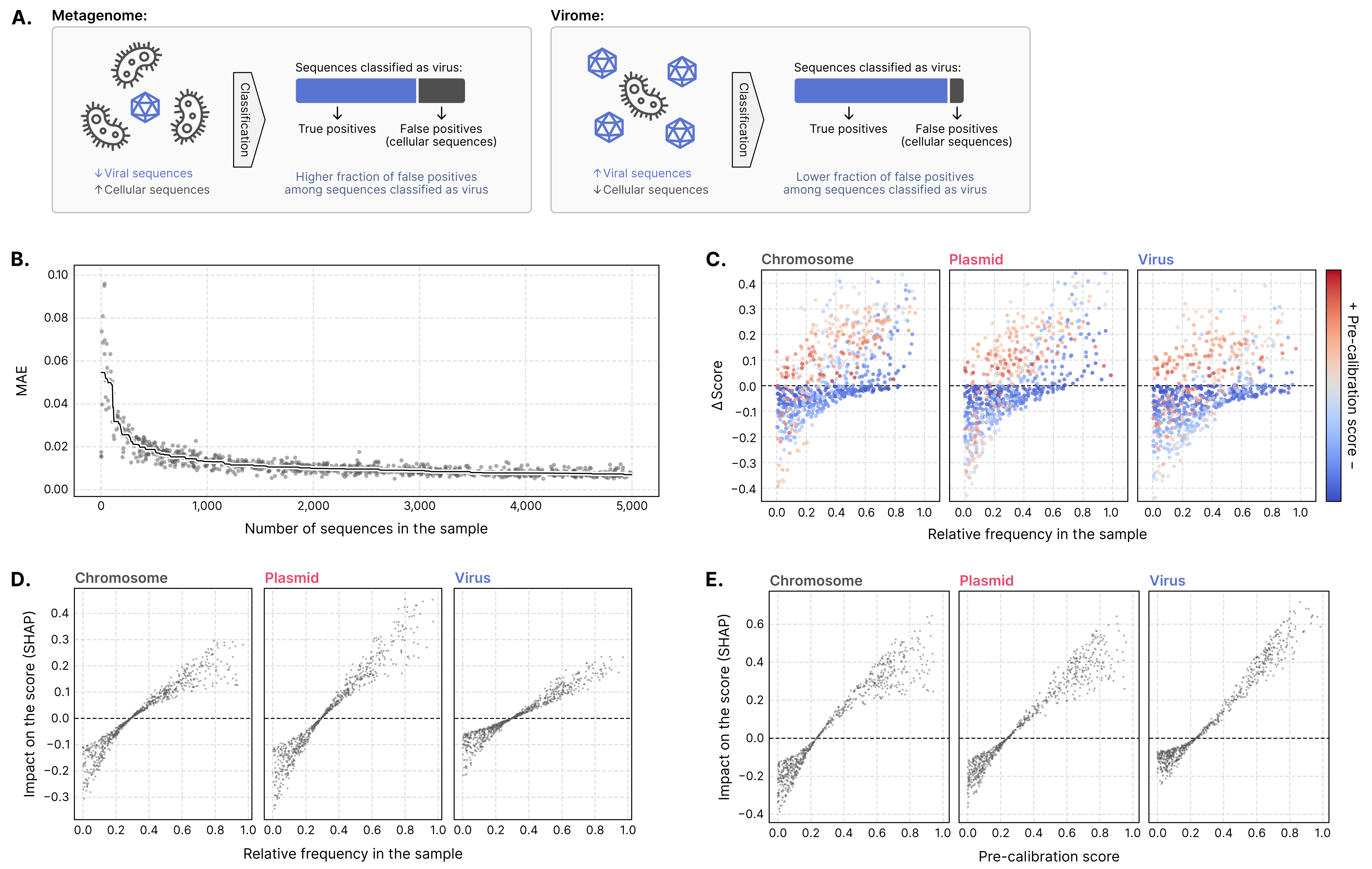

### Supplementary Figure 3

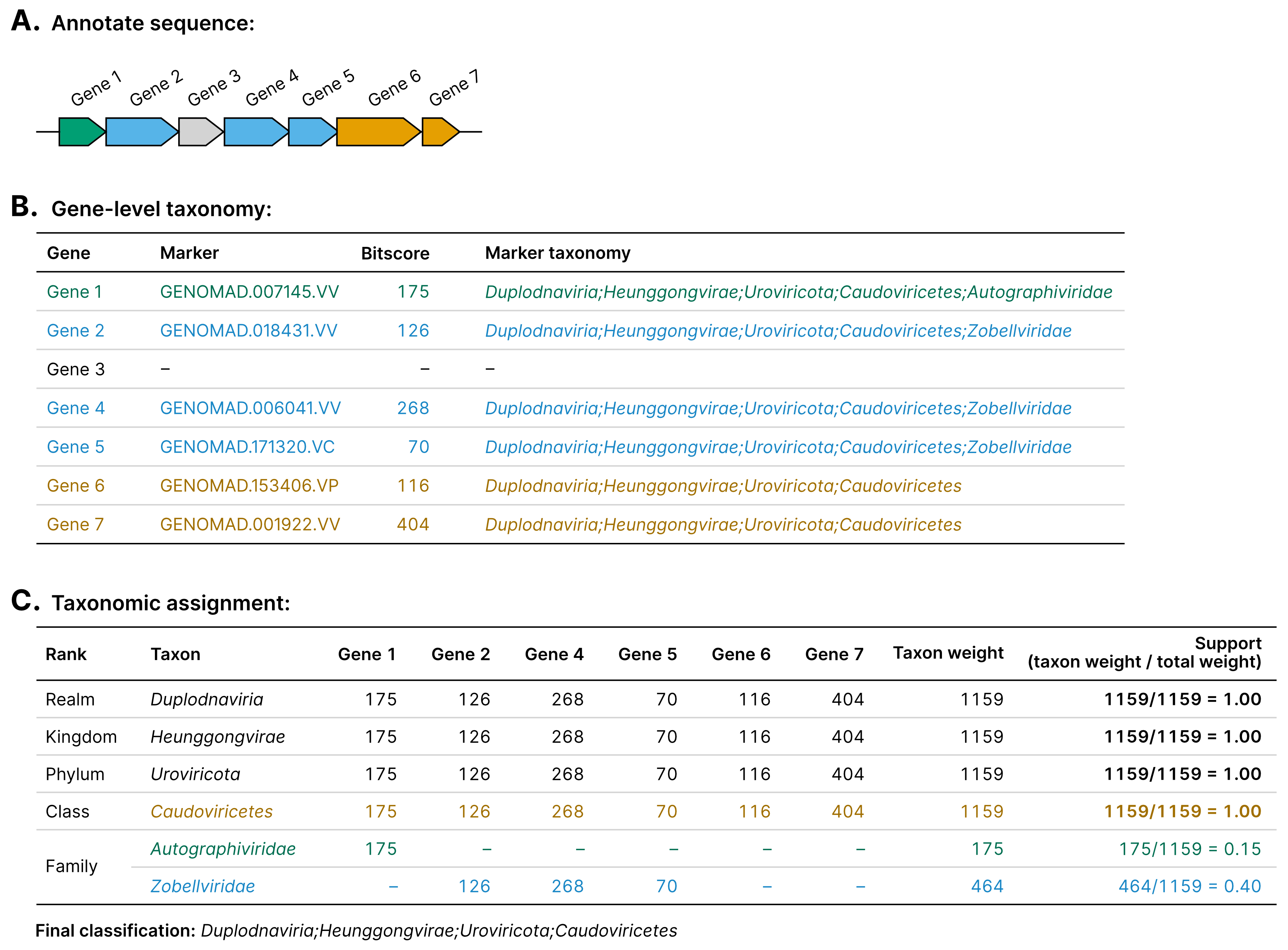

### Supplementary Figure 4

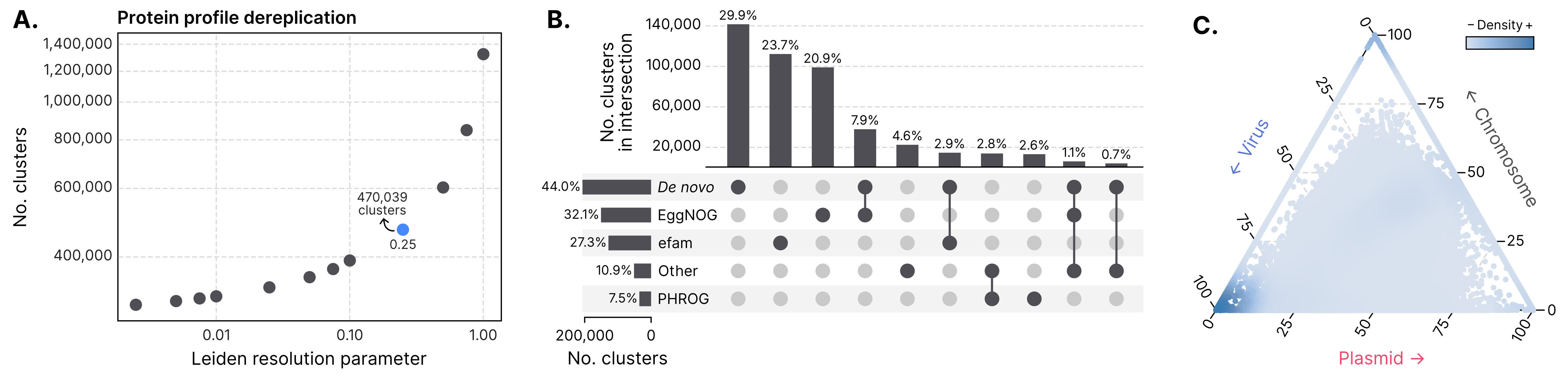

### Supplementary Figure 5

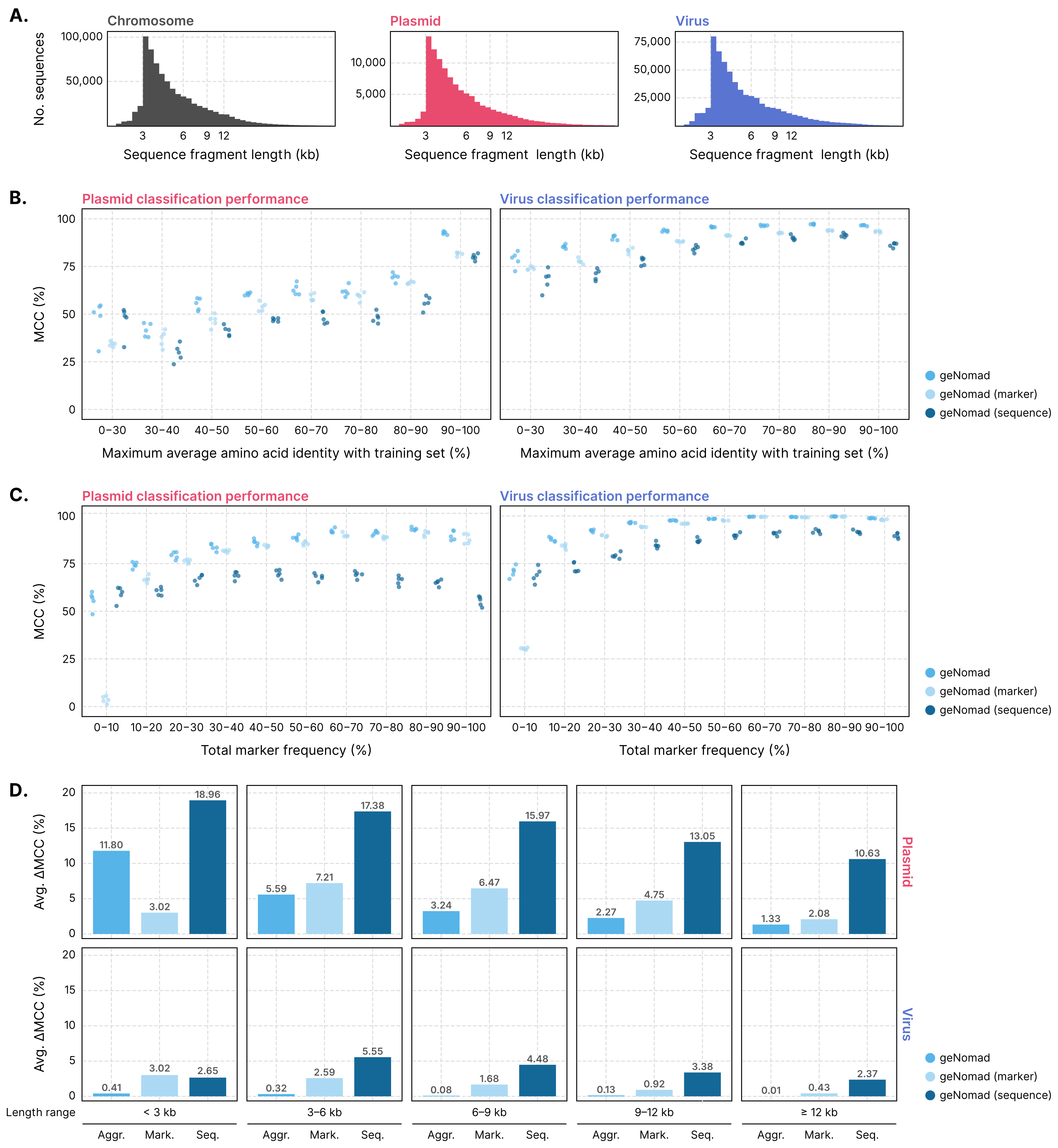

### Supplementary Figure 6

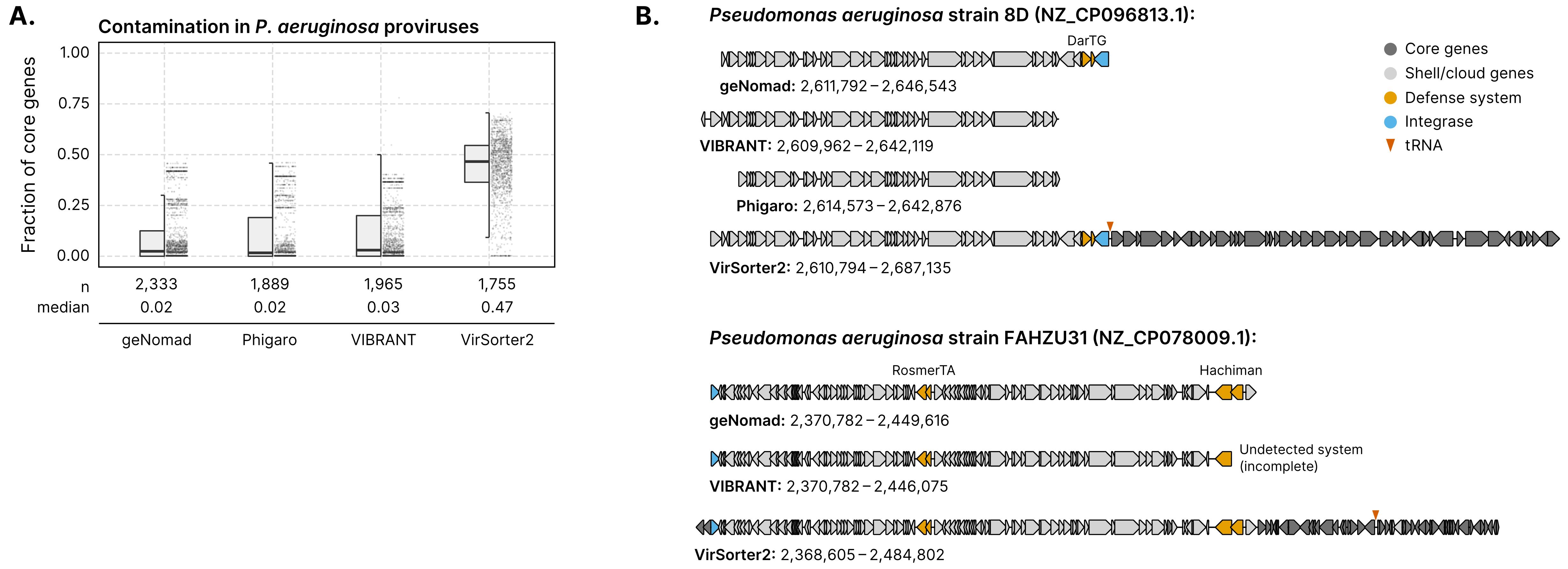
